# Extraordinary selection on the human X chromosome associated with archaic admixture

**DOI:** 10.1101/2022.09.19.508556

**Authors:** L. Skov, M. Coll Macià, E. Lucotte, M.I.A. Cavassim, D. Castellano, M.H. Schierup, K. Munch

## Abstract

The X chromosome in non-African human populations shows less diversity and less Neanderthal introgression than expected under the standard neutral model. We analyzed 162 X chromosomes from human males worldwide and discovered 14 chromosomal regions where haplotypes of several hundred kilobases rapidly rose to high frequencies in non-Africans. These observations cannot be explained by neutral genetic drift in realistic demographic scenarios and are only consistent with partial selective sweeps produced by strong selection. Using an approach for inferring individual Neanderthal-derived haplotypes, which do not rely on an archaic reference genome, we further discover that the swept haplotypes are devoid of the archaic ancestry otherwise typical of the affected chromosomal regions. The ancient Ust’-Ishim male carries its expected proportion of these haplotypes, implying that the sweeps must have occurred between 45,000 and 55,000 years ago. Finally, we find that the chromosomal positions of sweeps overlap previously reported hotspots of selection in great ape evolution. We propose that this puzzling combination of observations points to a general mechanism of positive selection unique to the X chromosome.

## Introduction

Mammalian X chromosomes display extraordinary evolution compared to autosomes and a disproportionate effect on male fertility and genetic incompatibilities between species. They are enriched for genes expressed in testis and undergo specific silencing during male meiosis. In *Mus musculus* and *Bos taurus,* there is evidence of a tug-of-war between the X and Y chromosome (Larson et al. 2016; Rathje et al. 2019; Hughes et al. 2020). In these species, the intra-genomic conflict is driven by the dynamic co-amplification of Y and X-linked homologs termed ampliconic genes and is expected to accelerate sex-chromosome evolution and the accumulation of incompatibilities between emerging species. Reported evidence shows that the X chromosome was repeatedly subject to strong positive selection in the great apes. This selection is revealed as a loss of diversity in large regions of the X-chromosome (Nam et al. 2015). The regions targeted by selection in each ape species overlap and further tend to fall inside larger regions repeatedly targeted by selection in the ancestral species of humans and chimpanzees (Dutheil et al. 2015) and of humans and orangutans (K. Munch, unpublished). Together, these findings lead to the hypothesis that meiotic drive plays a prominent role in primate X chromosome evolution (Nam et al. 2015).

Additionally, the patterns of archaic human introgression on the X and Y chromosomes are different from those observed on the autosomes (Schierup 2020). More than 250,000 years ago, modern humans admixed into Neanderthals replacing their Y chromosome (Petr et al.). On this occasion, Neanderthals also received at least as much modern human admixture on their X chromosome as they did on the autosomes (Hubisz et al. 2020). In the more recent meeting in the middle east about 55,000 years ago, the direction of admixture was reversed and qualitatively different. Here the Neanderthal Y chromosome did not introgress into the out-of-Africa modern humans, and the Neanderthal X chromosome introgressed to a much smaller extent than its autosomes (Sankararaman et al. 2016). Today, several megabase-long sections of the X chromosome are depleted of Neanderthal introgression. These sections overlap the regions of the X chromosome repeatedly targeted by natural selection in the great apes, as discussed above.

Here we investigate whether the reduced nucleotide diversity in regions of the human X chromosome is due to positive selection. We find that large regions of the X chromosome, which total more than 17Mb, contain haplotypes shared by all non-African populations that rapidly rose to high frequency. To our surprise, all these haplotypes are entirely devoid of archaic introgression. We then speculate which selective mechanism could cause this puzzling combination of observations.

## Results

### Megabase-long high-frequency haplotypes are common in non-African X chromosomes

We analyzed high-coverage genomes across the globe from the Simons Genome Diversity Project (SGDP) (Mallick et al. 2016). We first surveyed the nucleotide diversity of each population in 200 kb windows along the X chromosome and compared it to the diversity on a representative autosome with a similar length, chromosome 7. Figures 1A and 1B show the diversity patterns for a representative set of African and non-African populations, located as shown in Figure 1C. Figure 1D shows the distribution of pairwise differences in these 200kb windows (Figures S1 and S2 show diversity in all populations). Whereas African populations show a relatively even amount of diversity across the X chromosome, similar to what we observe along a representative autosome, all populations outside Africa display megabase-sized regions with extremely reduced diversity, often in similar parts on the X chromosome. Similar instances of such low diversity are not seen on the representative autosome, indicating that such extreme depressions of diversity is a unique property of the X chromosome.

**Figure 1:**
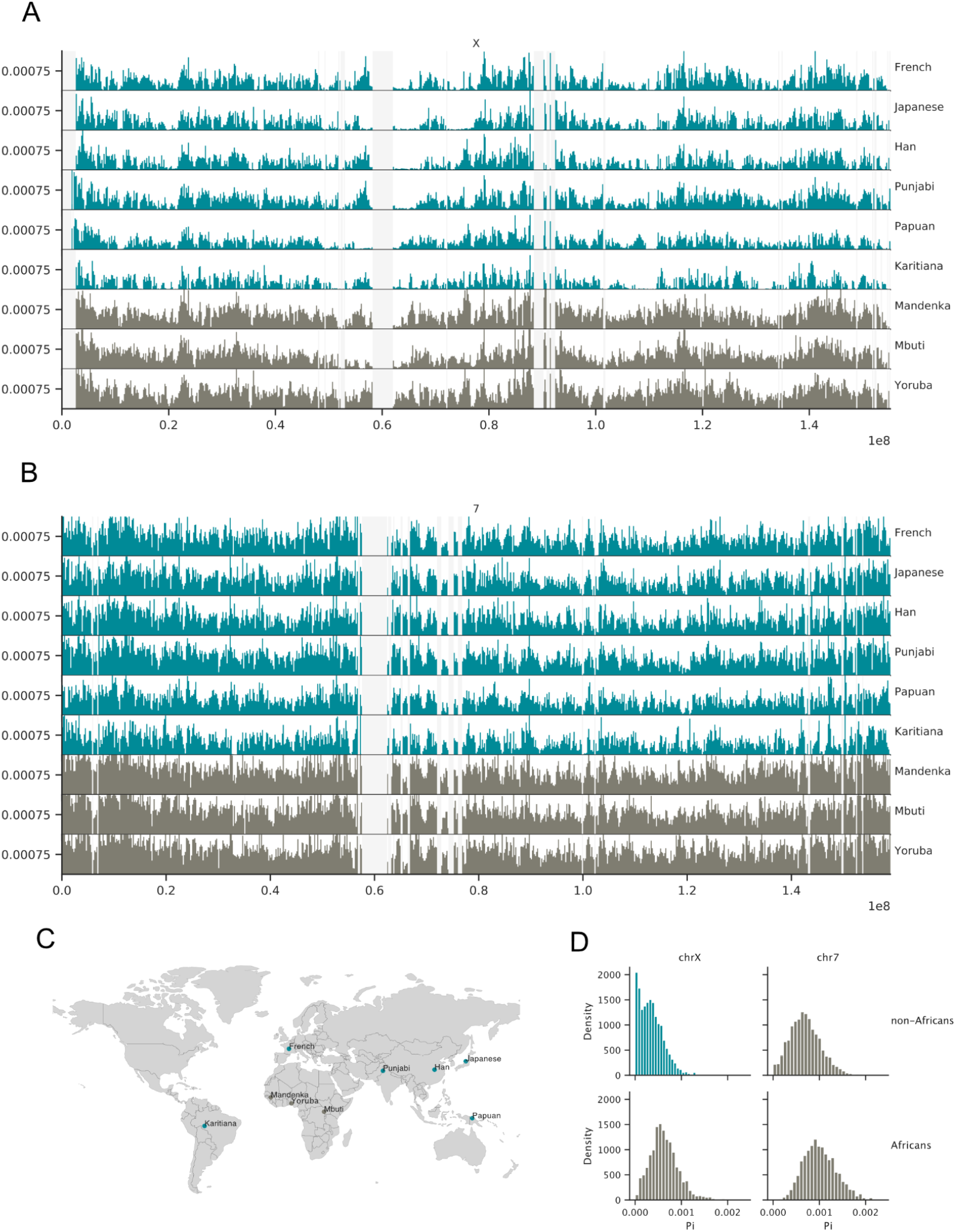
Population diversity across chromosomes X and 7. A) and B) Mean pairwise differences in 200kb windows across chromosomes X and 7 for populations from a set of representative populations where at least four X chromosomes are sampled. For a better visual comparison, the y-axis is truncated at 0.0015 to remove outliers. non-African and African populations are colored green and brown, respectively. Light grey regions represent missing data. C) Sampling origin of the selected populations. D) distribution of values in A and B divided into African or non-African populations.

**Figure 2:**
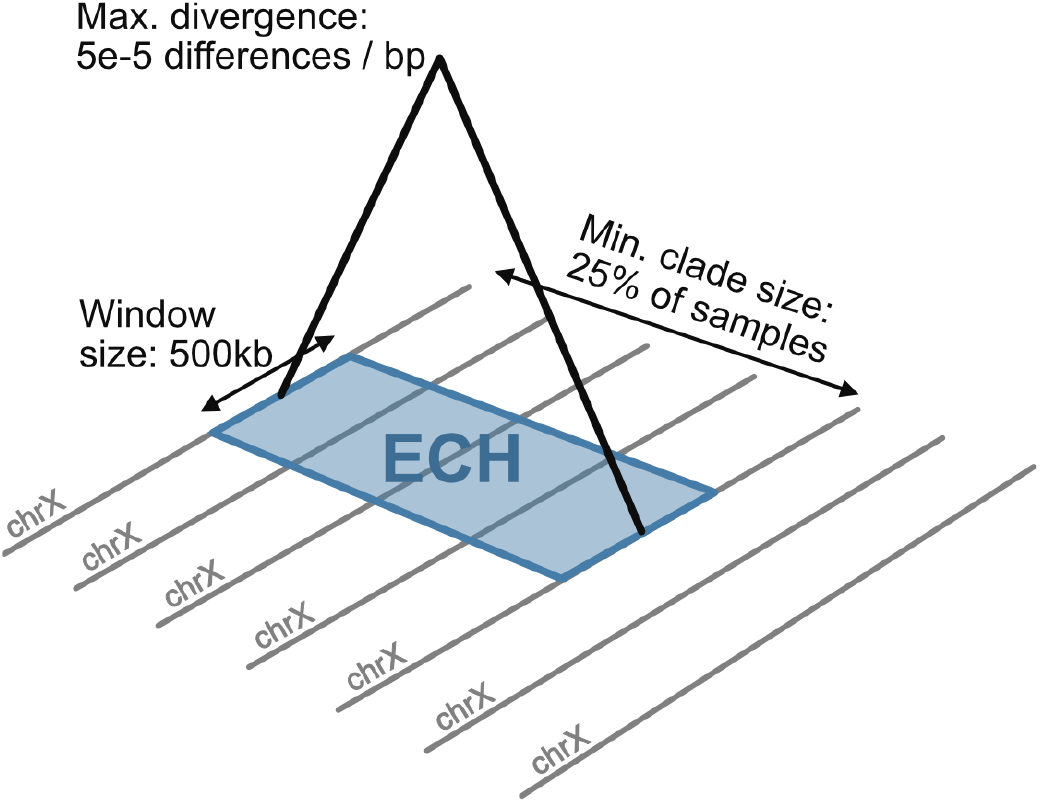
Definition of an extended common haplotype (ECH). Graphical depiction of the criteria used to define an ECH.

These initial observations suggest that high-frequency haplotypes are shared across non-African populations. To identify such haplotypes, we restricted the subsequent analysis to males for which the X chromosome is haploid. Following (Lucotte et al. 2018), we excluded males with missing data and males not showing the XY karyotype. We further removed African males with any evidence of recent European admixture (Supplementary Methods). This filtering left us with 162 males, of which 140 are non-Africans. (Supplementary Table S1). Sampling locations for the populations analyzed are shown in Figure 3A.

**Figure 3:**
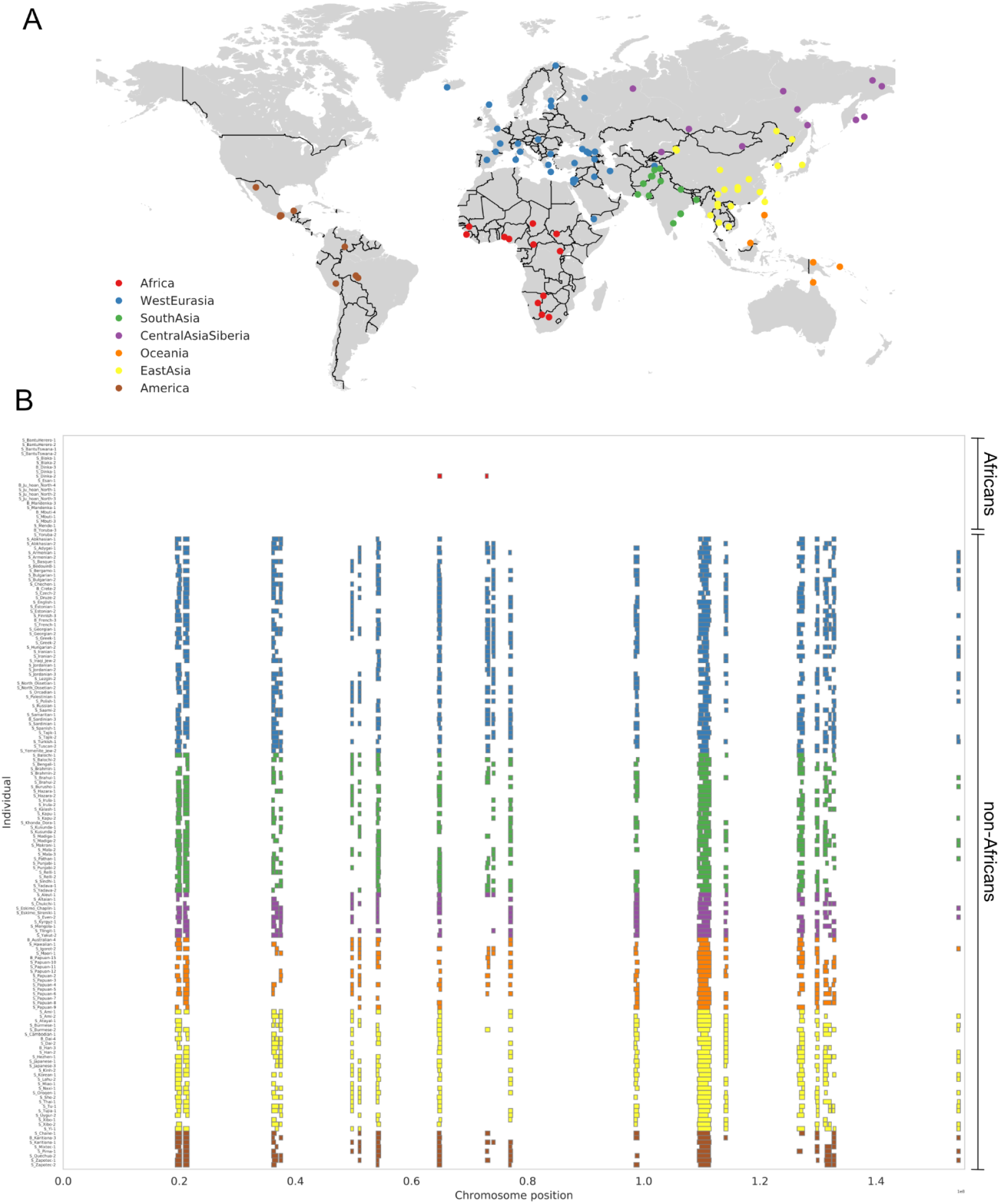
ECHs identified on each male X chromosome: A) World map showing sample locations. Colors represent the geographical region of each sample. B) ECHs on the X chromosome of each sampled male. Colors represent the geographical region of each sample as in A, with African samples shown at the top.

Next, we screened these X chromosomes for clades of long haplotypes (>500 kb) where the maximum genetic divergence between all member haplotypes is at most 0.005%. This divergence threshold corresponds to an expected common ancestry no more than 60,000 years ago (51,119 - 67,059, 50% probability mass), i.e., such clades should have a most recent common ancestor which is more recent than the out-of-Africa event (Mallick et al. 2016) (Supplementary Methods). When such haplotype clades contain at least 25% of the individuals in our data set, we refer to them as Extended Common Haplotypes (ECH) (See Figures S5 and S6 for the effect of alternative minimum clade sizes). We identified clades of ECHs in sliding windows of 500kb with a step size of 100kb (Figure 2) using a clique-finding algorithm (Bron and Kerbosch 1973). The ECHs identified in each individual are shown in Figure 3B. They are almost exclusively found in non-African populations with no apparent geographical differentiation in frequency outside of Africa. The sharing of haplotypes among non-Africans indicates that they rose to high frequency after the main exodus from Africa but before the subsequent diversification of non-African populations. In five of the nineteen chromosomal regions, the ECHs are shared by more than 75% of the non-African males and in fourteen by more than 50% of males. We will refer to this latter set as the fourteen most extreme regions. See Table S2 for hg19 coordinates of all ECH regions. The combined length of all nineteen ECHs is 17.3 Mb or 11% of the entire X chromosome. The ECHs are characterized by a higher proportion of high-frequency derived variants, which is expected if ECHs arose from a single initial haplotype (Figure 4).

**Figure 4:**
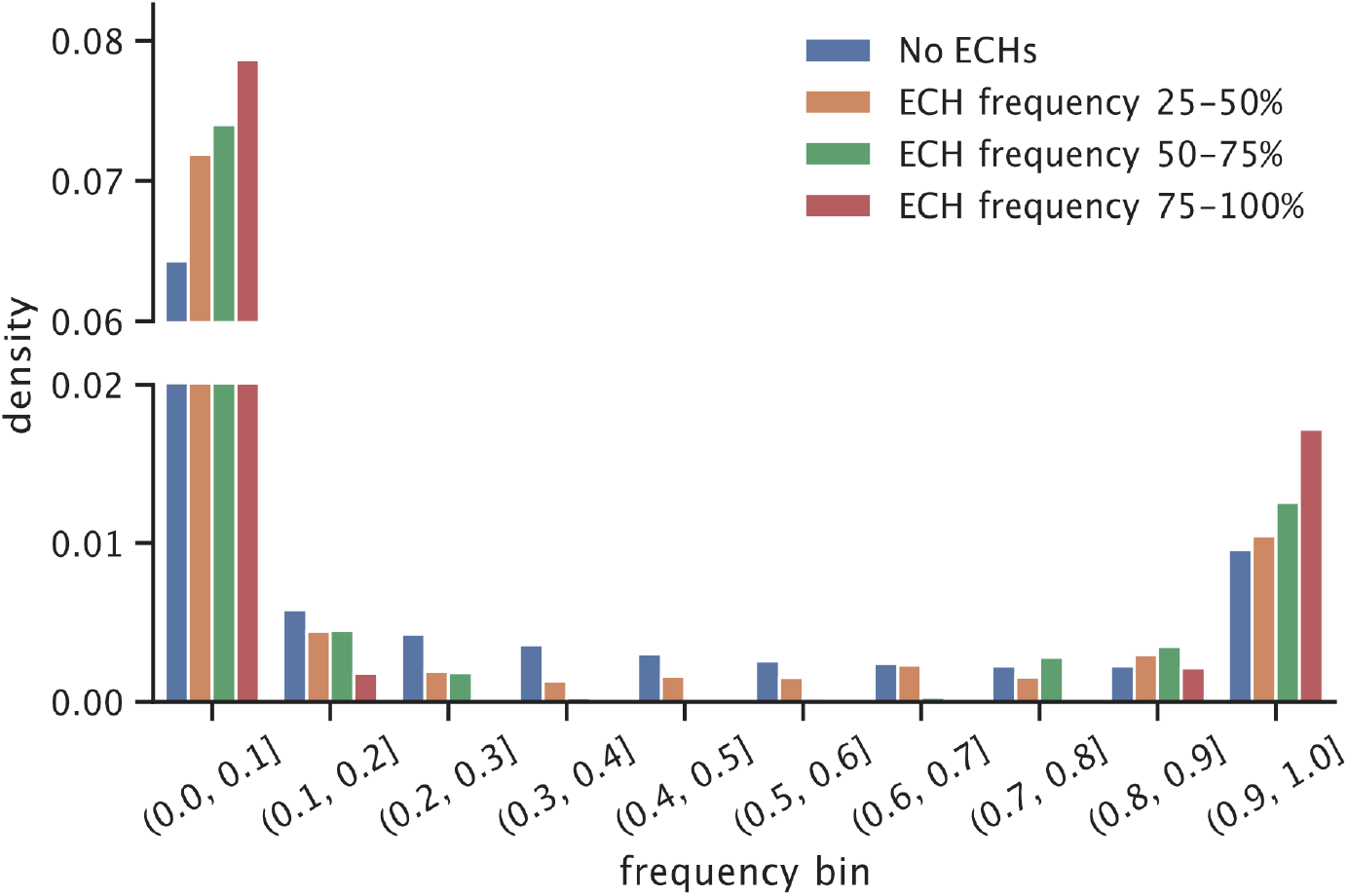
Site frequency spectrum and ECH frequency: Unfolded site frequency spectrum in four bins of ECH frequency. The y-axis is cut to accommodate a large number of rare variants.

Next, we visualized the haplotype structure of the chromosomal regions where each ECH is found and compared it to regions without ECHs. Figure 5A shows the core 900 kb region of the most extended and most frequent ECH with non-reference SNPs marked as black tics and the individual haploid X chromosomes clustered by UPGMA. All non-African X chromosomes form a single clade with highly reduced diversity compared to the African X-chromosomes, implying that a single ancestral haplotype rose to high frequency in non-Africans. In this example, the extended ECH haplotype spans at least 1.8Mb. The haplotype structure of the remaining ECHs shows the same pattern with one haplotype (and in some cases, two) shared by a large subset of non-African X chromosomes (Supplementary Data S1). For comparison, Figure 5B shows a typical 900kb region where we find no ECH. Here, non-African haplotypes do not form a single clade as more African X chromosome diversity is represented in non-Africans.

**Figure 5:**
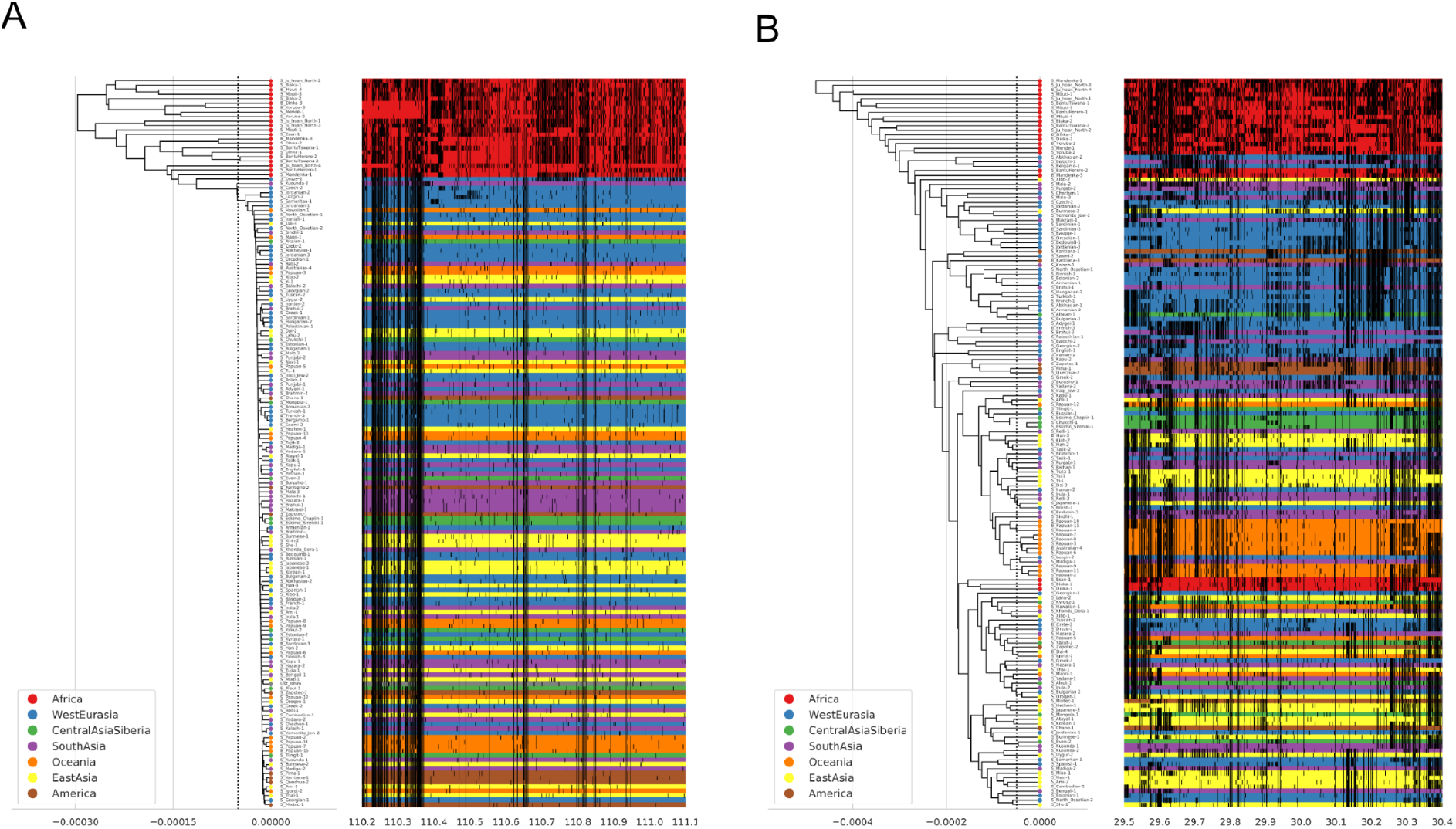
Examples of the relationship among haplotypes in identified regions: A) shows the core 900kb region around the longest and most frequent ECH (where the ECH frequency is at least 90% of its peak value) (Materials and Methods and Table S2). The left side of each figure is a UPGMA tree with the x-axis representing sequence divergence. The individual haplotypes are shown as horizontal lines on the right with the x-axis in hg19 megabase coordinates. Haplotypes are color-coded according to geographical region. The ancient Ust’-Ishim individual is marked with grey. Vertical black bars on each haplotype represents non-reference SNPs. B) For comparison, a 900kb region where no ECHs were identified.

Each individual carries many ECHs; on average, each non-African carries the ECH in 9.7 of the 14 regions where the ECH frequency is >50% (2.5 and 97.5 percentiles: 6 and 13). Figures 6 and S4 show the frequency of ECHs among non-Africans across the X chromosome and the core regions of each ECH shared by most individuals.

**Figure 6:**
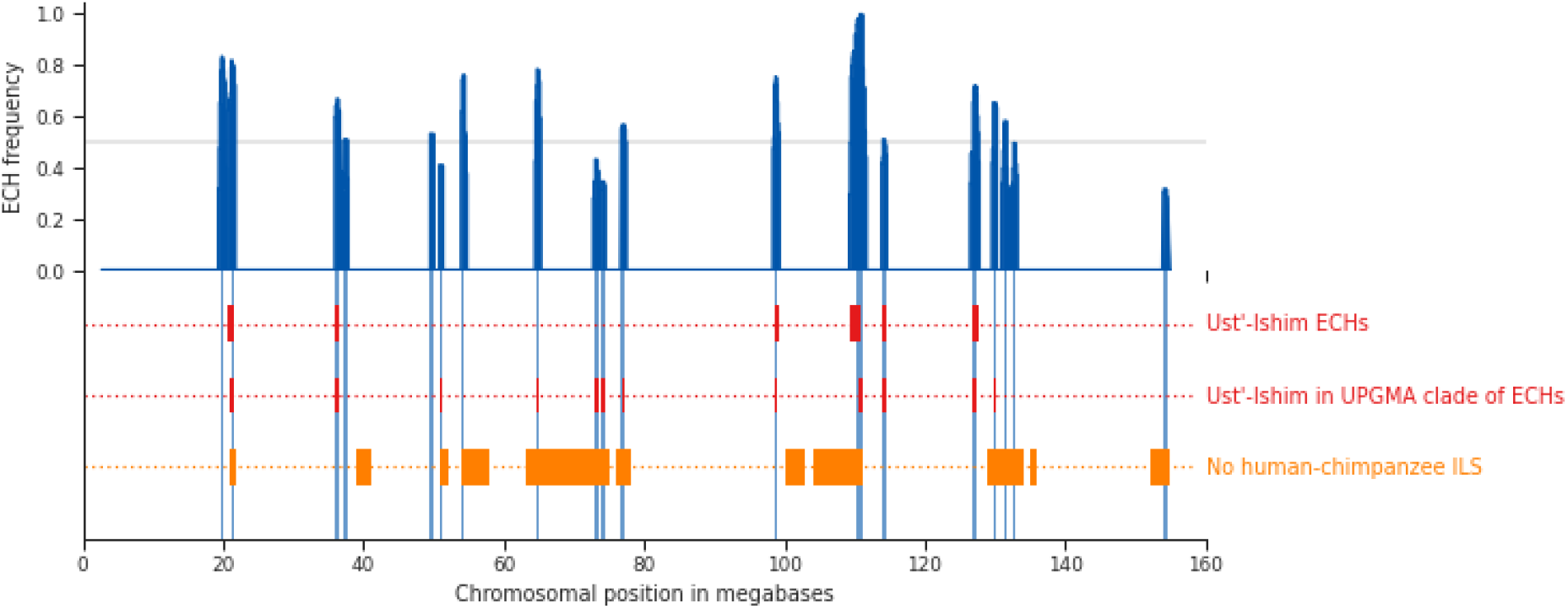
Frequency of ECHs along the X chromosome: Proportion of non-African haplotypes called as ECH in each 100kb window across the complete X chromosome (blue). In 14 cases, more than 50% of individual haplotypes are called as ECH. Vertical blue bands in lower each show the core regions around each peak where the proportion of non-Africans called as ECH is at least 90% of its peak value. ECHs in the Ust’-Ishim male, called using the clique finding approach, are shown in red. Regions depleted of incomplete lineage sorting between humans, chimpanzees, and gorillas are shown in orange.

### The rise in ECH frequency predated the Ust’-Ishim male

We included the ancient Ust’-Ishim genome dated at 45,000 BP (46,880-43,210 BP) (Fu et al. 2014a) to estimate further when these frequency changes occurred. The Ust’-Ishim is equally related to Europeans and Asians, suggesting that its lineage split off before the European/Asian population split, and it did not contribute directly to present-day diversity (Fu et al. 2014a). If haplotypes rose to high frequency soon after the main out-of-Africa event, we would expect the Ust’-Ishim individual to carry a number of ECHs that is similar to present-day non-Africans. To investigate this, we repeated our ECH-finding procedure, including the Ust’-Ishim male. The Ust’-Ishim male shared six ECHs of the 14 most frequent ECHs (Supplementary Methods). As an alternative approach, robust to false-positive SNP calls in the ancient sequence, we added the Ust’-Ishim male to haplotype plots of 500kb windows centered at each peak in Figure 6 (Supplementary Methods). The Ust’-Ishim falls inside a cluster of ECHs in 9 of the 14 most extreme regions, close to the non-African mean of 9.7 (Supplementary Methods). This similar number of ECHs carried by the Ust’-Ishim implies that most of the ECHs had already risen to high frequencies 45,000 years ago.

### Estimating the probability of observing the ECHs without natural selection

Our observations suggest that each ECH rose to high frequency from a single haplotype after the out-of-Africa event (60,000-80,000 BP) but before Ust’-Ishim lived (45,000 BP). This rapid change in frequency of such long haplotypes is striking and is characteristic of strong positive selection. However, genetic drift, particularly during extreme population bottlenecks, can also cause the frequency of long haplotypes to increase, and X chromosomes are more affected by bottlenecks than autosomes because of their smaller effective population size (Ne).

First, we performed simple frequency simulations of genetic drift to assess the probability of observing the haplotypes under neutral evolution. We simulated frequency trajectories to assess the probability that one 500kb haplotype across the entire chromosome rose to a high frequency by genetic drift in the span of time between the 45,000 BP of the Ust’-Ishim and the maximum ECH divergence of 60,000 years (Supplementary Methods). To cover a range of scenarios, we perform simulations for time spans of 5,000; 10,000; 15,000, 20,000; and 25,000 years (using additional evidence, we show later in the paper that only spans of 5,000 to 15,000 years are relevant). To find an appropriate bottleneck Ne for our simulations, we estimated a population demography, using SMC++, of the non-African samples used in our analysis (Supplementary Methods). Here we estimate the smallest autosomal effective population size (Ne) as 4,125. In our simulations, we conservatively assume an autosomal Ne of 3,000 and include, for comparison, a more unlikely autosomal Ne of 1,500. We perform simulations for two different X/autosome ratios of effective population size: One of 0.65 corresponds to the median ratio among African individuals in our data set, and a lower one of 0.51 corresponds to the median ratio of the non-Africans. The latter estimate may be smaller due to male-driven migration but may also be depressed by the removal of diversity in the ECHs, as discussed above. For an autosomal Ne of 3,000, these two X/autosome ratios correspond to X chromosomal Nes of 1950 and 975 (as shown in the legend for Figure 7). We further assume the sex averaged recombination rate of the X chromosome (1.16e-8) (Kong et al. 2010). We perform 500,000 simulations for each combination of parameters.

**Figure 7:**
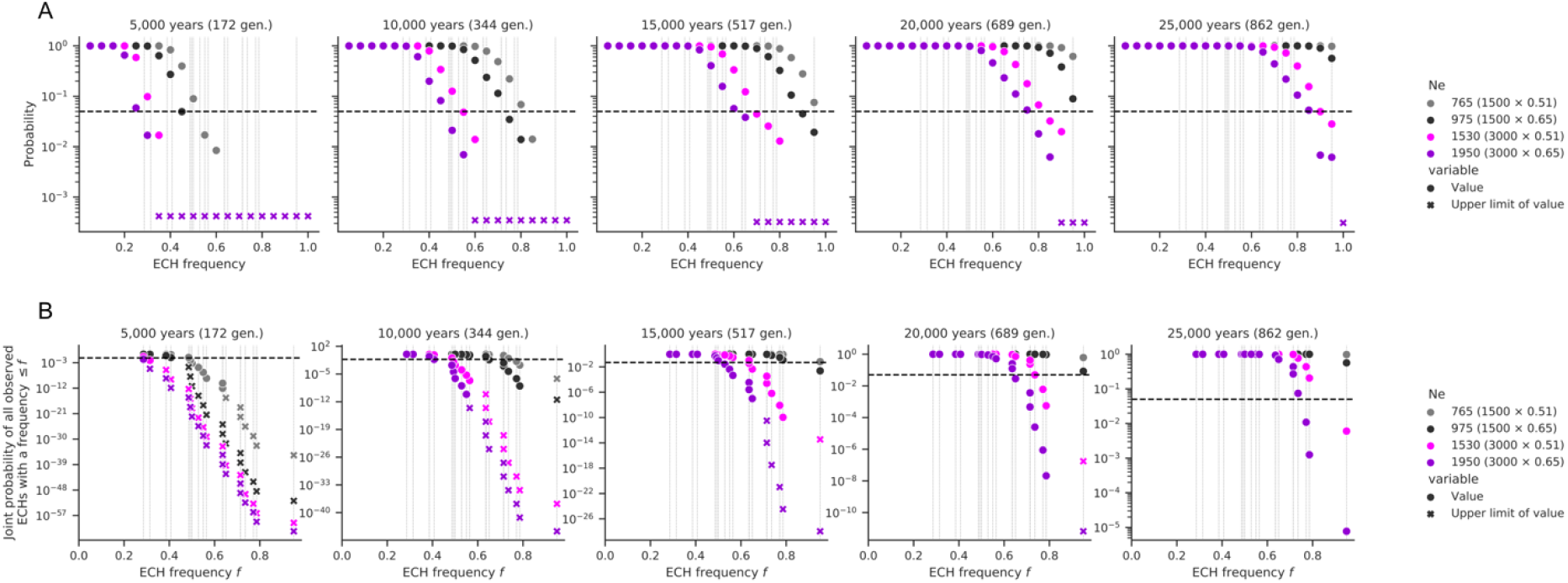
Estimated probability of ECHs: **A)** Circles represent evenly spaced probabilities as the fraction of simulations in which a 500kb haplotype along the chromosome rises to the specified or greater frequency in 500,000 simulations. Multiple scenarios are shown in each panel in which the duration of the bottleneck increases from left (5000 years) to the right (25,000 years) in intervals of 5,000 years. The legend lists the X chromosome population sizes used and shows the autosomal population size and X/autosome ratio in parentheses. Simulations assume a mean recombination rate of 1.16e-8 and a generation time of 29 years. Colors represent different combinations of autosomal Ne and X/autosome ratio combinations, as shown in the legend. Crosses represent upper bounds on the probability in each case where no haplotypes reached the target frequency in the simulation. A dashed horizontal line marks the 0.05 value. Thin vertical lines represent the frequencies of the ECHs we identify in the SGDP male data set. **B)** Same as A) but represents the joint probability of the identified 500kb ECHs below a given frequency. In each facet, the leftmost point thus represents the probability of observing only the lowest frequency ECH, and the rightmost point shows the joint probability of observing all the ECHs we identify.

Simulations are summarized in Figure 7, where panel A presents the probability that at least one 500kb haplotype along the chromosome rises to the given frequency inside the given span of time. Assuming realistic parameters, a 5,000-15,000 year span, and an autosomal bottleneck population size of 3,000, we find that while drift can create a few low-frequency ECHs, the probabilities of creating each high-frequency ECHs are extremely low. Figure 7B quantifies the joint probability of observing all the identified ECHs at the frequencies they each occur. Each point represents the joint probability of the observed ECH with that frequency together with all observed ECHs at a lower frequency. The rightmost points in each facet thus show the joint probability of all observed ECHs. For the parameter range stated above, the observed ECHs are extremely unlikely to have arisen neutrally. Grey and black points show the probabilities for the autosomal population size of 1500. Even with this unrealistically low autosomal population size, the time span inferred from Ust-ishim admixture also needs to be unrealistically long (20,000-25,000 years) for the joint probability of all observed ECHs to exceed 0.05. This shows that neutral processes alone are very unlikely to have caused the observed ECHs.

Second, we performed forward simulations of full X chromosomes using a population demography and a recombination map. Since the SGDP data results from a very extensive population structure and demographics not well suited for simulation, we chose the homogeneous CEU population (Utah residents with Northern and Western European ancestry) from the 1000 genomes data set as our model population. We repeated our inference procedure on the CEU population and then performed forward simulations to assess the probability that the ECHs called in this population were produced by genetic drift.

In our inference of ECHs on the 49 CEU male X chromosomes, we need to consider that a large sample from this single homogeneous population will much more often find recent common ancestry than the highly structured SGDP. To accommodate this contribution to the clade size of such early coalescences, we increase the ECH clade size from 25%, used in our main analysis, to 50%. Inference using this clade size reproduces 15 of the 19 ECH peaks identified among males from the SGDP. In the CEU population, the ECHs together cover 10% of the entire chromosome, similar to the 11% in the SGDP data set. In the subsequent simulations of the past 200,000 generations, performed using SLiM3 (Haller and Messer 2019), we use the previously published population size trajectory for CEU that, to our knowledge, estimates the strongest bottleneck (Gravel et al. 2011). The simulations also use a fine-scale pedigree-based recombination map of the X chromosome (Kong et al. 2010). We perform simulations using the same X/autosome Ne ratios (0.51 and 0.65) used in our frequency simulations. Across 500 simulations of 49 male X chromosomes, we compute the analyzed proportion of the X chromosome where an ECH is called, and the analyzed proportion of 100kb sequence windows called as ECH across all simulated chromosomes. Assuming the African X/autosome ratio of 0.65, these two proportions are 0.3% (0 - 2.0%) and 0.2% (0 - 1.2%) (interval is 2.5 and 97.5 percentiles across whole chrX simulations). Assuming instead a lower non-African X/autosome ratio of 0.51 (which would be lower in part due to the sweeps), the numbers are 1.1% (0 - 4.0%) and 0.7% (0 - 2.5%). In our analysis of the 49 CEU samples, these two proportions are a magnitude larger (11.1% and 7.6%). Considering further that these CEU proportions are 2.8 and 3.0 times larger than the 97.5 percentile of proportions obtained from simulations (Figure 8), it is extremely unlikely that the ECHs arose neutrally in the CEU population. It is even less likely that they arose neutrally in only 10,000 years, as revealed by the analysis of the SGDP male data set.

**Figure 8:**
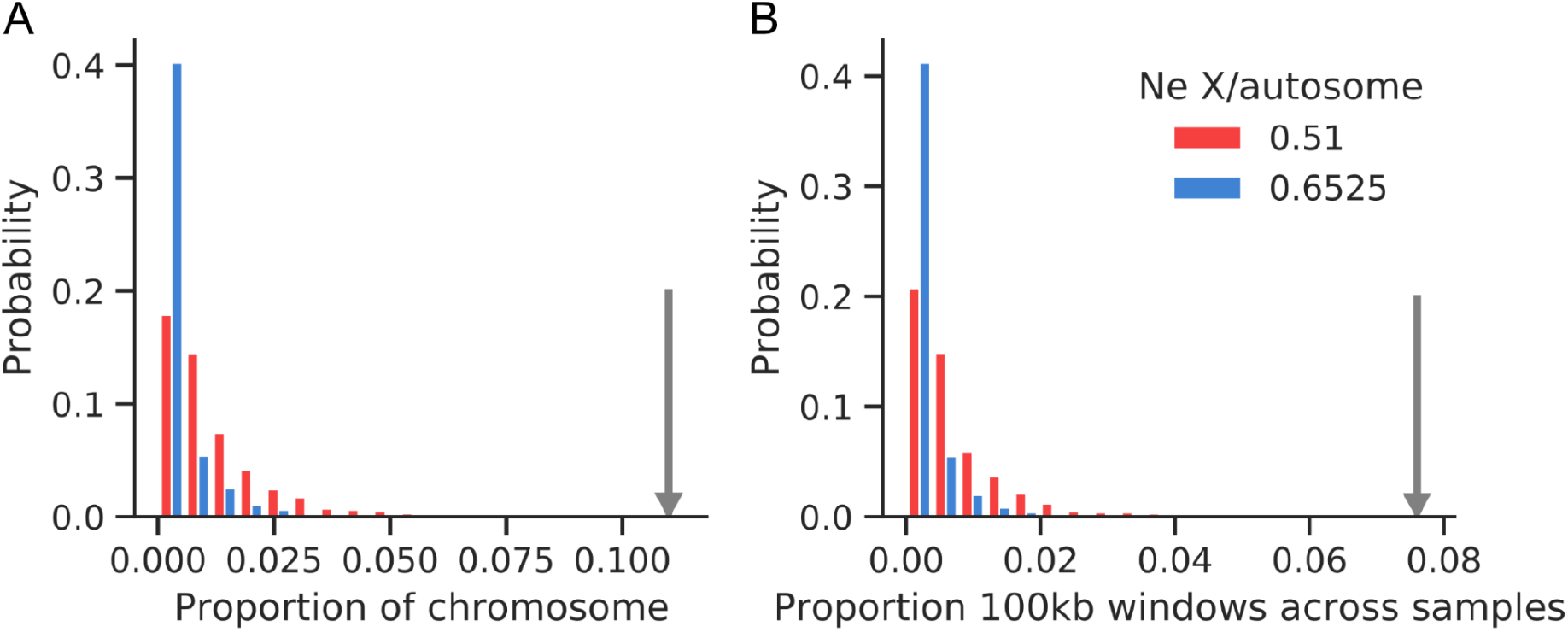
Simulated proportions of ECHs on the X chromosome in the CEU population: 500 forward simulations (SLiM3) of 49 male X chromosomes using a CEU population size demography and a parent-offspring recombination map. **A)** The proportion of the X chromosome where an ECH is called. Colors represent the X to autosome population size ratio used in simulations. The gray arrow shows the value of this statistic in the CEU population. **B)** As for A, but showing the proportion of 100kb sequence windows called as ECH across all simulated sample sequences.

These analyses indicate that positive selection must have driven the ECHs to high frequencies.

### Selective sweeps on the X chromosome may be recurrent

We have previously reported evidence of selective sweeps in the human-chimpanzee ancestor (Dutheil et al. 2015) and Munch unpublished (Supplementary methods). We, therefore, tested if these overlap with the independent observation of ECH regions we report here. We find a strong overlap between the ECHs and the sections of the chromosome swept at least once during the 2-4 million years that separated the human-chimpanzee and human-gorilla speciation events (Scally et al. 2012; Munch et al. 2016), shown as orange blocks in Figure 6 (Jaccard stat.: 0.13, p-value: 1.6e-4 (Supplementary Methods).

### Selective sweeps displaced archaic introgressed sequence on the X chromosome

The admixture with archaic humans that followed the main out-of-Africa event left a far smaller proportion of introgressed sequence in the X chromosome than in the autosomes. In non-African populations, the Neanderthal component is thus only 0.3% compared to 1.4% for the autosome. Denisovan admixture is virtually absent on the X chromosome, with only Oceanians carrying a small proportion (0.18%) (Sankararaman et al. 2016). To investigate a relationship between archaic admixture and the selective sweeps we detect, we applied the approach of (Skov et al. 2018) to call genomic segments of archaic human ancestry in each non-African male X chromosome (Supplementary Methods). This approach uses a hidden Markov model to search for clusters of derived SNPs not observed in an un-admixed group of African genomes. Compared to inference in a previous report (Sankararaman et al. 2016), the method we apply can identify archaic segments not represented in sequenced archaic genomes. This approach estimates a mean admixture proportion across individuals of 0.8%. However, when we restrict to archaic segments which share derived variants with the high coverage archaic genomes (Denisova, Vindija, and Altai), we identify a proportion of introgressed sequence (0.36%) similar to that previously reported (Supplementary Methods and Table S3).

Restricting the analysis to the 100kb windows where some individual carries an ECH (11% of the X chromosome), we observe that the archaic proportion here is 0.3%, compared to 0.9% in windows where no individuals carry an ECH (t-test p-value 6e-26). To investigate if this reduction is caused by low levels of admixture in ECHs specifically, we first extracted the subset of chromosomal 100kb windows where any individuals carry an ECH. In each of these 100kb windows, we computed the mean archaic admixture proportion of the ECHs and of the haplotypes in the same positions that are not part of the ECH clade. We find that the ECHs are almost completely without inferred archaic admixture. In contrast, the remaining haplotypes in the same genomic windows have a mean admixture proportion of 0.70%, ranging between 0.35% and 1.03%, depending on the geographical region. This admixture proportion is close to the archaic contribution in chromosomal regions not overlapping an ECH. The analysis thus reveals that the absence of admixture in the ECHs themselves entirely explains the reduced archaic admixture in the chromosomal regions where they appear. Within the ECHs, the mean proportion of archaic admixture is 0.0045%, corresponding to a reduction of 99% compared to the non-ECH haplotypes in the same chromosomal windows. This proportion is consistent with a complete absence of archaic admixture since our admixture inference is associated with a small false-positive rate (Skov et al. 2018). Each geographical region displays this absence of archaic admixture in ECHs (Figure 9A), and so does each core ECH region of the chromosome where archaic admixture is detected (Figure 9B). If archaic admixture contributed significantly to diversity, we might falsely conclude that ECHs were admixture-free simply because the elevated diversity was not compatible our definition of ECHs. However, we rule out that the calling of ECHs is affected by the presence/absence of admixture by performing a supplementary analysis where we mask admixture segments identified in each individual before calling ECHs (Supplementary Methods, Figure S8).

**Figure 9:**
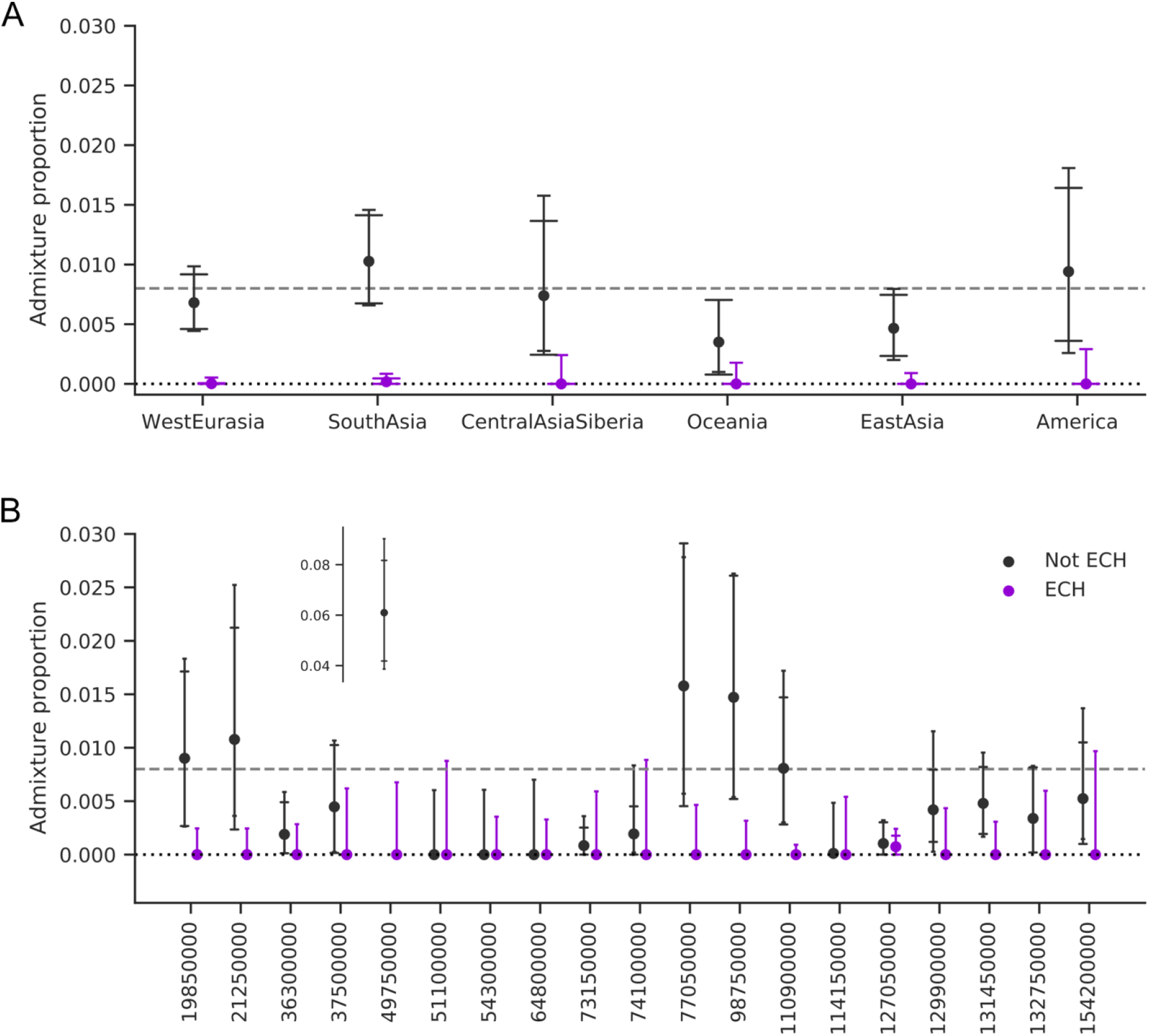
Admixture proportions in chromosomal regions of partial sweeps. Mean admixture proportions of ECHs (purple) and in remaining haplotypes (black) are computed separately for each 100kb window. Error bars with wide caps designate the standard 95% confidence intervals obtained from 10,000 bootstrapping iterations. Error bars with narrow caps designate Jeffrey’s binomial confidence interval, which better represents very low frequencies (in computing this interval, we assume that the sample size equals the number of 100kb sequence windows). The dotted line represents zero admixture. The dashed line shows the mean admixture proportion on the X chromosome. A) Mean admixture proportions of haplotypes from each geographical region. B) Admixture proportions at each individual ECH. X-axis labels represent chromosomal positions where each ECH has the highest frequency (peaks in Figure 6). For legibility, one outlier is shown separately.

### Archaic displacement provides further support for positive selection

While EHCs displaced archaic admixture, their rise in frequency cannot be a result of purifying selection against archaic contribution. While such negative selection has most likely occurred (Harris and Nielsen 2016) it cannot explain why a single admixture-free haplotype, rather than many, would rapidly rise in frequency. In contrast, this would be expected if a single haplotype was under positive selection for reasons unrelated to archaic admixture. In this context, archaic admixture thus merely serves as a backdrop against which selection on admixture-free ECHs is visible.

Our finding that ECHs also displaced Neanderthal admixture in the Ust’-Ishim, which lived 45,000 BP (Fu et al. 2014b), further allows us to narrow the period in which the ECHs rose in frequency, as the sweeps must necessarily have happened between the admixture event and the Ust’-Ishim. From the length of Neanderthal admixture segments in the Ust’-Ishim male, the main Neanderthal admixture event has been dated to 9599 years before the Ust’-Ishim. Using four standard errors as the confidence interval on this estimate implies that the sweeps must have occurred in a span of time that is at least 3,724 years and at most 15,341 years (Supplememtary methods). This implies that the time intervals of 20,000 and 25,000 years included in our simulations (Figure 7) fall outside the range of relevant parameters, further decreasing the probability that ECHs arose from neutral processes.

## Discussion

Our analyses suggest that nineteen regions of the X chromosome, totaling 11% of its length, carry long high-frequency haplotypes (ECHs) that are shared across all non-African populations. The frequency of these haplotypes rose after the out-of-Africa (OoA) event and the subsequent archaic admixture events. These dramatic frequency changes appear to have been fully or almost fully completed by the time of the Ust-Ishim, dated to 45,000 BP. The large size of these haplotypes is consistent with a rapid increase in frequency not compatible with a neutral process of genetic drift. We conclude that they must have increased in frequency by positive selection. Surprisingly, the ECHs are entirely free of archaic admixture, suggesting that whatever variants drove their rise in frequency, these arose on haplotypes without archaic admixture.

Fourteen of the identified ECHs each span between 500kb and 1.8Mb in at least 50% of non-Africans (Table S2). The strongest sweep spans 900kb in 91% of non-Africans and affects 53% of non-Africans across a 1.8Mb region. In comparison, the strongest selective sweep reported from human diversity data is at the lactase gene and spans 800kb in 77% of European Americans (Bersaglieri et al. 2004). The selection coefficient on the focal SNP has been estimated to be 1.6-1.8% (Mathieson and Mathieson 2018; Stern et al. 2019), suggesting that the selection coefficients responsible for several ECHs may have been well above 1%.

We have not identified specific genetic elements associated with the ECH rise in frequency. The ECHs strongly overlap regions depleted of ILS in the common ancestor of humans and chimpanzees, suggesting that they underwent repeated selection in great ape evolution. This raises the possibility that our observations reflect processes repeatedly affecting the X chromosome across evolutionary timescales.

We have previously suggested a role of ampliconic genes that show postmeiotic expression in mouse testis (Mueller et al. 2008, 2013) and are involved in sex chromosomal meiotic drive processes in mouse (Cocquet et al. 2012; Larson et al. 2016) and fruit files (Ellison and Bachtrog 2019). However, while human ampliconic regions are significantly proximal to the swept regions (permutation test, p-value: 0.024), they do not generally overlap. The core regions of each ECH, shared by most individuals, each include several genes, and we do not detect enrichment for any gene ontology (Supplementary Methods). Protein-coding genes also show no enrichment of genes with elevated expression in testis (Supplementary Methods). One sweep, however, only has a single protein-coding gene, ACTRT1, at its center, which is linked to spermatid formation (Heid et al. 2002).

At the present time, we cannot envision a likely scenario that explains all our observations. We cautiously hypothesize that the selective sweeps may be due to sex chromosome meiotic drive: If an averagely even transmission of X and Y-sperm in meiosis is maintained by a dynamic equilibrium of antagonizing drivers on X and Y, the bottlenecked main out-of-Africa population may have been invaded by sex chromosome drivers retained in an earlier out-of-Africa population. This hypothesis is indirectly supported by recent evidence suggesting that the rapid expansion of the FT Y-chromosome haplotype originated in East/Southeast Asia 50-55,000 BP and displaced Y lineages carried by the later main wave out of Africa (Hallast, Agdzhoyan, et al. 2020). The spread of a Y-driver across Eurasia would be followed by X haplotypes spreading from the same source population to restore an even meiotic transmission in the populations invaded by Y-drivers. The date of the Ust’-Ishim male, which carries this FT haplotype, would place these events in the 45-55,000 BP window where we conclude the sweeps occurred. An Asian origin of EHCs and subsequent displacement of the main wave out of Africa is also consistent with our observation that ECHs displaced not only Neanderthal admixture in west Eurasia but also the Denisovan admixture in Asia. If this hypothesis is true, the swept regions represent the only remaining haplotypes from early non-African populations not admixed with Neanderthals. This hypothesis provides testable predictions that will guide our future work.

Whatever the explanation, we believe it must be unique to X chromosomes, possibly in the form of other consequences to fertility or the fidelity of meiosis. We thus suggest that future studies could focus on surveying any male fertility consequences of the ECHs that we report, perhaps in combination with specific Y chromosome haplogroups. Large cohorts with male fertility data and genome-wide sequencing or genotyping will soon be available for such a study.

## References

Bersaglieri, T., Sabeti, P.C., Patterson, N., Vanderploeg, T., Schaffner, S.F., Drake, J.A., Rhodes, M., Reich, D.E., and Hirschhorn, J.N. (2004). Genetic signatures of strong recent positive selection at the lactase gene. American Journal of Human Genetics 74, 1111–1120. https://doi.org/10.1086/421051.

Bron, C., and Kerbosch, J. (1973). Algorithm 457: finding all cliques of an undirected graph. Commun Acm 16, 575–577. https://doi.org/10.1145/362342.362367.

Cocquet, J., Ellis, P.J., Mahadevaiah, S.K., Affara, N.A., Vaiman, D., and Burgoyne, P.S. (2012). A Genetic Basis for a Postmeiotic X Versus Y Chromosome Intragenomic Conflict in the Mouse. PLoS Genetics 8. https://doi.org/10.1371/journal.pgen.1002900.

Dutheil, J.Y., Munch, K., Nam, K., Mailund, T., and Schierup, M.H. (2015). Strong Selective Sweeps on the X Chromosome in the Human-Chimpanzee Ancestor Explain Its Low Divergence. PLOS Genetics 11, e1005451. https://doi.org/10.1371/journal.pgen.1005451.

Ellison, C., and Bachtrog, D. (2019). Recurrent gene co-amplification on Drosophila X and Y chromosomes. Plos Genet 15, e1008251. https://doi.org/10.1371/journal.pgen.1008251.

Fu, Q., Li, H., Moorjani, P., Jay, F., Slepchenko, S.M., Bondarev, A.A., Johnson, P.L., Aximu-Petri, A., Prüfer, K., Filippo, C. de, et al. (2014a). Genome sequence of a 45,000-year-old modern human from western Siberia. Nature 514, 445–449. https://doi.org/10.1038/nature13810.

Fu, Q., Li, H., Moorjani, P., Jay, F., Slepchenko, S.M., Bondarev, A.A., Johnson, P.L.F., Aximu-Petri, A., Prüfer, K., Filippo, C. de, et al. (2014b). Genome sequence of a 45,000-year-old modern human from western Siberia. Nature 514, 445. https://doi.org/10.1038/nature13810.

Gravel, S., Henn, B.M., Gutenkunst, R.N., Indap, A.R., Marth, G.T., Clark, A.G., Yu, F., Gibbs, R.A., Bustamante, C.D., Altshuler, D.L., et al. (2011). Demographic history and rare allele sharing among human populations. Proc National Acad Sci 108, 11983–11988. https://doi.org/10.1073/pnas.1019276108.

Harris, K., and Nielsen, R. (2016). The Genetic Cost of Neanderthal Introgression. Genetics 203, 881–891. https://doi.org/10.1534/genetics.116.186890.

Heid, H.W., Figge, U., Winter, S., Kuhn, C., Zimbelmann, R., and Franke, W.W. (2002). Novel Actin-Related Proteins Arp-T1 and Arp-T2 as Components of the Cytoskeletal Calyx of the Mammalian Sperm Head. Exp Cell Res 279, 177–187. https://doi.org/10.1006/excr.2002.5603.

Hubisz, M.J., Williams, A.L., and Siepel, A. (2020). Mapping gene flow between ancient hominins through demography-aware inference of the ancestral recombination graph. Plos Genet 16, e1008895. https://doi.org/10.1371/journal.pgen.1008895.

Hughes, J.F., Skaletsky, H., Pyntikova, T., Koutseva, N., Raudsepp, T., Brown, L.G., Bellott, D.W., Cho, T.-J., Dugan-Rocha, S., Khan, Z., et al. (2020). Sequence analysis in Bos taurus reveals pervasiveness of X-Y arms races in mammalian lineages. Genome Res https://doi.org/10.1101/gr.269902.120.

Kong, A., Thorleifsson, G., Gudbjartsson, D.F., Masson, G., Sigurdsson, A., Jonasdottir, A., Walters, G.B., Jonasdottir, A., Gylfason, A., Kristinsson, K.Th., et al. (2010). Fine-scale recombination rate differences between sexes, populations and individuals. Nature 467. https://doi.org/10.1038/nature09525.

Larson, E.L., Keeble, S., Vanderpool, D., Dean, M.D., and Good, J.M. (2016). The composite regulatory basis of the large X-effect in mouse speciation. Molecular Biology and Evolution msw243. https://doi.org/10.1093/molbev/msw243.

Lucotte, E.A., Skov, L., Jensen, J.M., Macià, M.C., Munch, K., and Schierup, M.H. (2018). Dynamic Copy Number Evolution of X- and Y-Linked Ampliconic Genes in Human Populations. Genetics 209, 907–920. https://doi.org/10.1534/genetics.118.300826.

Mallick, S., Li, H., Lipson, M., Mathieson, I., Gymrek, M., Racimo, F., Zhao, M., Chennagiri, N., Nordenfelt, S., Tandon, A., et al. (2016). The Simons Genome Diversity Project: 300 genomes from 142 diverse populations. Nature 538. https://doi.org/10.1038/nature18964.

Mathieson, S., and Mathieson, I. (2018). FADS1 and the Timing of Human Adaptation to Agriculture. Mol Biol Evol 35, 2957–2970. https://doi.org/10.1093/molbev/msy180.

Mueller, J.L., Mahadevaiah, S.K., Park, P.J., Warburton, P.E., Page, D.C., and Turner, J.M.A. (2008). The mouse X chromosome is enriched for multicopy testis genes showing postmeiotic expression. Nat Genet 40, 794–799. https://doi.org/10.1038/ng.126.

Mueller, J.L., Skaletsky, H., Brown, L.G., Zaghlul, S., Rock, S., Graves, T., Auger, K., Warren, W.C., Wilson, R.K., and Page, D.C. (2013). Independent specialization of the human and mouse X chromosomes for the male germ line. Nature Genetics https://doi.org/10.1038/ng.2705.

Munch, K., Nam, K., Schierup, M., and Mailund, T. (2016). Selective Sweeps across Twenty Millions Years of Primate Evolution. Molecular Biology and Evolution 33, msw199.https://doi.org/10.1093/molbev/msw199.

Nam, K., Munch, K., Hobolth, A., Dutheil, J., Veeramah, K.R., Woerner, A.E., Hammer, M.F., Project, G., Mailund, T., Schierup, M., et al. (2015). Extreme selective sweeps independently targeted the X chromosomes of the great apes. Proceedings of the National Academy of Sciences 112, 6413–6418. https://doi.org/10.1073/pnas.1419306112.

Petr, M., Hajdinjak, M., Fu, Q., Essel, E., Rougier, H., Crevecoeur, I., Semal, P., Golovanova, L.V., Doronichev, V.B., Lalueza-Fox, C., et al. (n.d.). The evolutionary history of Neanderthal and Denisovan Y chromosomes. Science (New York, N.Y.) 369, 1653–1656. https://doi.org/10.1126/science.abb6460.

Rathje, C., Johnson, E., Drage, D., Patinioti, C., Silvestri, G., Affara, N., Ialy-Radio, C., Cocquet, J., Skinner, B., and Ellis, P. (2019). Differential Sperm Motility Mediates the Sex Ratio Drive Shaping Mouse Sex Chromosome Evolution. Curr Biol 29, 3692–3698.e4. https://doi.org/10.1016/j.cub.2019.09.031.

Sankararaman, S., Mallick, S., Patterson, N., and Reich, D. (2016). The Combined Landscape of Denisovan and Neanderthal Ancestry in Present-Day Humans. Current Biology 26, 1241–1247. https://doi.org/10.1016/j.cub.2016.03.037.

Scally, A., Dutheil, J.Y., Hillier, L.W., Jordan, G.E., Goodhead, I., Herrero, J., Hobolth, A., Lappalainen, T., Mailund, T., Marques-Bonet, T., et al. (2012). Insights into hominid evolution from the gorilla genome sequence. Nature 483, 169–175. https://doi.org/10.1038/nature10842.

Schierup, M.H. (2020). The last pieces of a puzzling early meeting. Sci New York N Y 369, 1565–1566. https://doi.org/10.1126/science.abe2766.

Skov, L., Hui, R., Shchur, V., Hobolth, A., Scally, A., Schierup, M., and Durbin, R. (2018). Detecting archaic introgression using an unadmixed outgroup. Plos Genet 14, e1007641. https://doi.org/10.1371/journal.pgen.1007641.

Stern, A.J., Wilton, P.R., and Nielsen, R. (2019). An approximate full-likelihood method for inferring selection and allele frequency trajectories from DNA sequence data. PLOS Genetics 15, e1008384. https://doi.org/10.1371/journal.pgen.1008384.

Terhorst, J., Kamm, J.A., and Song, Y.S. (2017). Robust and scalable inference of population history from hundreds of unphased whole genomes. Nature Genetics 49, 303–309. https://doi.org/10.1038/ng.3748.

Bersaglieri T., P. C. Sabeti, N. Patterson, T. Vanderploeg, S. F. Schaffner, et al., 2004 Genetic signatures of strong recent positive selection at the lactase gene. American journal of human genetics 74: 1111–20. https://doi.org/10.1086/421051

Bron C., and J. Kerbosch, 1973 Algorithm 457: finding all cliques of an undirected graph. Commun Acm 16: 575–577. https://doi.org/10.1145/362342.362367

Cocquet J., P. J. Ellis, S. K. Mahadevaiah, N. A. Affara, D. Vaiman, et al., 2012 A Genetic Basis for a Postmeiotic X Versus Y Chromosome Intragenomic Conflict in the Mouse. PLoS Genetics 8. https://doi.org/10.1371/journal.pgen.1002900

Dutheil J. Y., K. Munch, K. Nam, T. Mailund, and M. H. Schierup, 2015 Strong Selective Sweeps on the X Chromosome in the Human-Chimpanzee Ancestor Explain Its Low Divergence. PLOS Genetics 11: e1005451. https://doi.org/10.1371/journal.pgen.1005451

Ellison C., and D. Bachtrog, 2019 Recurrent gene co-amplification on Drosophila X and Y chromosomes. Plos Genet 15: e1008251. https://doi.org/10.1371/journal.pgen.1008251

Fu Q., H. Li, P. Moorjani, F. Jay, S. M. Slepchenko, et al., 2014a Genome sequence of a 45,000-year-old modern human from western Siberia. Nature 514: 445. https://doi.org/10.1038/nature13810

Fu Q., H. Li, P. Moorjani, F. Jay, S. M. Slepchenko, et al., 2014b Genome sequence of a 45,000-year-old modern human from western Siberia. Nature 514: 445–449. https://doi.org/10.1038/nature13810

Haller B. C., and P. W. Messer, 2019 SLiM 3: Forward Genetic Simulations Beyond the Wright-Fisher Model. Mol Biol Evol 36: 632–637. https://doi.org/10.1093/molbev/msy228

Harris K., and R. Nielsen, 2016 The Genetic Cost of Neanderthal Introgression. Genetics 203: 881–891. https://doi.org/10.1534/genetics.116.186890

Heid H. W., U. Figge, S. Winter, C. Kuhn, R. Zimbelmann, et al., 2002 Novel Actin-Related Proteins Arp-T1 and Arp-T2 as Components of the Cytoskeletal Calyx of the Mammalian Sperm Head. Exp Cell Res 279: 177–187. https://doi.org/10.1006/excr.2002.5603

Hubisz M. J., A. L. Williams, and A. Siepel, 2020 Mapping gene flow between ancient hominins through demography-aware inference of the ancestral recombination graph. Plos Genet 16: e1008895. https://doi.org/10.1371/journal.pgen.1008895

Hughes J. F., H. Skaletsky, T. Pyntikova, N. Koutseva, T. Raudsepp, et al., 2020 Sequence analysis in Bos taurus reveals pervasiveness of X-Y arms races in mammalian lineages. Genome Res. https://doi.org/10.1101/gr.269902.120

Kong A., G. Thorleifsson, D. F. Gudbjartsson, G. Masson, A. Sigurdsson, et al., 2010 Fine-scale recombination rate differences between sexes, populations and individuals. Nature 467. https://doi.org/10.1038/nature09525

Larson E. L., S. Keeble, D. Vanderpool, M. D. Dean, and J. M. Good, 2016 The composite regulatory basis of the large X-effect in mouse speciation. Molecular biology and evolution msw243. https://doi.org/10.1093/molbev/msw243

Lucotte E. A., L. Skov, J. M. Jensen, M. C. Macià, K. Munch, et al., 2018 Dynamic Copy Number Evolution of X- and Y-Linked Ampliconic Genes in Human Populations. Genetics 209: 907–920. https://doi.org/10.1534/genetics.118.300826

Mallick S., H. Li, M. Lipson, I. Mathieson, M. Gymrek, et al., 2016 The Simons Genome Diversity Project: 300 genomes from 142 diverse populations. Nature 538. https://doi.org/10.1038/nature18964

Mathieson S., and I. Mathieson, 2018 FADS1 and the Timing of Human Adaptation to Agriculture. Mol Biol Evol 35: 2957–2970. https://doi.org/10.1093/molbev/msy180

Mueller J. L., S. K. Mahadevaiah, P. J. Park, P. E. Warburton, D. C. Page, et al., 2008 The mouse X chromosome is enriched for multicopy testis genes showing postmeiotic expression. Nat Genet 40: 794–799. https://doi.org/10.1038/ng.126

Mueller J. L., H. Skaletsky, L. G. Brown, S. Zaghlul, S. Rock, et al., 2013 Independent specialization of the human and mouse X chromosomes for the male germ line. Nature Genetics. https://doi.org/10.1038/ng.2705

Munch K., K. Nam, M. Schierup, and T. Mailund, 2016 Selective Sweeps across Twenty Millions Years of Primate Evolution. Molecular biology and evolution 33: msw199. https://doi.org/10.1093/molbev/msw199

Nam K., K. Munch, A. Hobolth, J. Dutheil, K. R. Veeramah, et al., 2015 Extreme selective sweeps independently targeted the X chromosomes of the great apes. Proceedings of the National Academy of Sciences 112: 6413–8. https://doi.org/10.1073/pnas.1419306112

Petr M., M. Hajdinjak, Q. Fu, E. Essel, H. Rougier, et al., The evolutionary history of Neanderthal and Denisovan Y chromosomes. Science (New York, N.Y.) 369: 1653–1656. https://doi.org/10.1126/science.abb6460

Rathje C., E. Johnson, D. Drage, C. Patinioti, G. Silvestri, et al., 2019 Differential Sperm Motility Mediates the Sex Ratio Drive Shaping Mouse Sex Chromosome Evolution. Curr Biol 29: 3692–3698.e4. https://doi.org/10.1016/j.cub.2019.09.031

Sankararaman S., S. Mallick, N. Patterson, and D. Reich, 2016 The Combined Landscape of Denisovan and Neanderthal Ancestry in Present-Day Humans. Current Biology 26: 1241–1247. https://doi.org/10.1016/j.cub.2016.03.037

Scally A., J. Y. Dutheil, L. W. Hillier, G. E. Jordan, I. Goodhead, et al., 2012 Insights into hominid evolution from the gorilla genome sequence. Nature 483: 169–75. https://doi.org/10.1038/nature10842

Schierup M. H., 2020 The last pieces of a puzzling early meeting. Sci New York N Y 369: 1565–1566. https://doi.org/10.1126/science.abe2766

Skov L., R. Hui, V. Shchur, A. Hobolth, A. Scally, et al., 2018 Detecting archaic introgression using an unadmixed outgroup. Plos Genet 14: e1007641. https://doi.org/10.1371/journal.pgen.1007641

Stern A. J., P. R. Wilton, and R. Nielsen, 2019 An approximate full-likelihood method for inferring selection and allele frequency trajectories from DNA sequence data. PLOS Genetics 15: e1008384. https://doi.org/10.1371/journal.pgen.1008384

