## Supplementary methods for "Extraordinary selection on the human X chromosome associated with archaic admixture"

#### Dataset

We analyzed male X chromosomes from the fully public subset of genomes published by the Simons Genome Diversity Project (Mallick et al. 2016). We the version of this data originally published in 2016. We exclude African individuals with evidence of gene flow from non-African populations: Mozabite and Saharawi show extensive non-African ancestry and Neanderthal admixture (Mallick et al. 2016), and Masai and Somali show a non-African component in STRUCTURE analysis (Mallick et al. 2016), Luhya and close populations Luo and BantuKenya, show a non-African component in Admixture analysis (Auton et al. 2015). Following (Mallick et al. 2016), we also exclude five samples (S_Finnish-1, S_Finnish-2, S_Mansi-1, S_Mansi-2, S_Palestinian-2) "based on missing X chromosome data in the initial processing, for themselves or a second sample." We further exclude S_Lezgin-1, for which sex is not assigned, and S_Palestinian-2 and S_Naxi-2, which show patterns of sequencing coverage not congruent with their assigned sex (Lucotte et al. 2018). This filtering leaves us with 239 individuals, of which 162 are males. We list these individuals in Table S1.

#### Visualization of diversity including both males and females

We performed an initial analysis of diversity across populations using both males and females. Figures S1 and S2 show the mean pairwise difference in 200kb windows across chromosomes X and 7 in each population. We discarded windows with fewer than 50,000 called positions (50%) and considered these windows missing data. For a better visual comparison, the y-axis is truncated at 0.0015 to remove outliers. Grey regions represent missing data.

#### Pairwise sequence differences on chrX in males

We initially remove the two pseudoautosomal regions of chromosome X because these recombine with the Y chromosome. We then computed the proportion of pairwise differences along the chromosome for each pair of male individuals in non-overlapping windows of 100kb. We discarded windows with fewer than 50,000 called positions (50%) and considered these missing data.

#### Inference of admixture segments

We use a hidden Markov model to infer archaic segments (Skov et al. 2018). All scripts for using the method are available online at the GitHub repository: https://github.com/LauritsSkov/Introgression-detection. The method finds archaic genomic segments in non-Africans by identifying segments with a strong enrichment of single nucleotide variants that are not seen in an unadmixed outgroup population (in this case, Sub-Saharan populations). Non-African variants not present in the outgroup show a ten-fold enrichment in archaic segments because these variants have been accumulated since the common ancestor of archaic and modern humans.

We use an outgroup consisting of individuals from two datasets. We use all Sub-Saharan Africans (populations: YRI, MSL, ESN) from the 1000 Genomes Project that does not have a component shared with the European population (Auton et al. 2015) figure 2a) and all Sub-Saharan African populations from SGDP except Masai, Somali, Sharawi, and Mozabite ((Mallick et al. 2016) see supporting figure 8.1), which show signs of out-of-Africa admixture. For all data, we removed sites that fell within repeat-masked regions (downloaded from the UCSC genome browser) as well as sites that were not in the strict callability mask for the 1000 Genomes Project: (ftp://ftp.1000genomes.ebi.ac.uk/vol1/ftp/release/20130502/supporting/accessible_genome_masks/StrictMask/).

Since the method is sensitive to variation in mutation rate, we calculate the background mutation rate using the variant density of all variants from populations YRI, LWK, GWD, MSL, and ESN in windows of 100 Kb divided by the mean variant density of the whole genome.

The hidden Markov model was trained using the whole genome and not only the X chromosome. The reason for this is that there are not enough archaic segments to accurately infer transition and emission parameters just using the X chromosome. To accommodate the smaller effective population size of the X chromosome, we scaled the emission values using the following approach: Since males contribute 3/4 of mutations, but the X chromosome only spends a third of its time in males, then the X chromosome mutation rate is 5/6=0.8333 of the autosomal one, i.e., if the autosomal emission of state 2 is 0.38, it should be 0.3166 for X.

We show the effect of decoding with haploid X chromosome parameters versus decoding with autosomal parameters in Table S3. This table also lists the proportions of archaic admixture in autosomes. Not relying on the sequence of the admixing archaic human allows this method to predict admixture not represented by the diversity of sequenced archaic genomes. However, when we condition that fragments share derived alleles with the archaic individual (Altai, Vindija, or Denisova), we obtain admixture proportions similar to those of (Sankararaman et al. 2016).

Archaic segments are identified as regions where the posterior probability of the archaic state in the HMM is at least 0.8. Segments interleaved by more than 25Mb of missing data are split in two. 1kb windows with less than 200 bases called are excluded. We then compute the mean proportion of archaic sequence in each non-overlapping 100kb window. Windows, where over 20,000 bases are not called, are treated as missing data.

Because several swept regions show a local reduction in Ne, it is important to note that this does not impede our ability to detect admixture. The power of our inference method relies on the fact that the density of SNPs private to non-Africans is low compared to the density of SNPs in archaic segments. For this reason, reduced effective population size in the regions where we identify ECHs, will increase rather than decrease our power to detect admixture.

#### Identification of Extended Common Haplotypes (ECHs)

We compute the pairwise distance between all the male X chromosomes in sliding windows of 500kb with a step of 100kb. For each window, we identify sets of haplotypes where the pairwise distance between all pairs is smaller than 5e-5. These sets must include at least 25% of individuals in the dataset (at least 40 males). We do this by constructing a graph where nodes represent individuals, and edges connect individuals with a distance smaller than 5e-5. We then use a clique-finding algorithm (Bron and Kerbosch 1973) to extract fully connected sets of at least 40 individuals. We refer to each haplotype in such a set as an Extended Common Haplotype (ECH). An ECH may be longer than 500kb if it spans several overlapping 500kb windows. Figures S4 and S5 show the distribution of ECHs called for alternative clique sizes of 32 and 49 (20% and 30% of individuals, respectively).

The maximal pairwise distance of 5e-5 corresponds to an expected common ancestry of 59,520 years BP and was chosen to postdate the out-of-Africa event. The date is obtained assuming a mutation rate of 4.2e-10 = 4.3e-10 * 0.8 * 1.221. 4.3e-10 is the autosomal mutation rate (Fu et al. 2014), 0.8 corrects for sex-specific mutation rate and hemizygosity of the X, and 1.221 corrects for a contribution of false-positive SNPs in the SGDP data set. The latter correction is computed as the ratio between divergence to the human reference of sample S_Eskimo_Sireniki-1 at two filtering levels provided in the SGDP data set: filtering level 9 (used in this study and recommended for population genetic analysis) and the strictest filtering level 1 (used for estimation of mutation rate) (Mallick et al. 2016).

As described above, 100kb sequence windows with fewer than 50,000 called positions (50%) are considered missing data. Chromosomal 100kb, where this is true for more than 10% of individuals are masked, leaving 138,5 Mb of analyzed chromosome in the downstream analyses.

#### Defining chromosomal regions with peak proportions of ECHs

In each chromosomal region where we identify ECHs, we use a peak detection algorithm (Du, Kibbe et al. 2006) to find the chromosomal position where most individuals carry the ECH and identify from that the consecutive 100kb windows that share this maximal number of individuals. We report the coordinates of these peak regions as "peak start" and "peak end" in Table S2. In addition, we operationally define a region around each peak where the proportion of individuals called as ECH is at least 90% of the peak value (Figure S3). Similarly, we define a wider region where the proportion of individuals called as ECH rises above 75% of the peak value. Table S2 lists chromosomal coordinates of both 90%-regions and 75%-regions along with the minimum proportion of non-African haplotypes called as ECH across each region, as well as the proportion of haplotypes that span each entire region.

#### Data quality in ECH and non-ECH windows

We find that ECH windows have fewer missing bases than other windows: The proportion of missing bases in sequence windows called as ECH is 23% compared to 31% in windows we do not call as ECH. As described above, we discard 100kb windows of individiuals where more than 50% of bases are missing. This implies that pairwise distances to individuals with missing data in a 100kb window cannot be computed. Chromosomal windows where ECHs are called have fewer missing pairwise distances than the remaining windows: In ECH windows, the proportion of missing pairwise distances is 0.009%, whereas it is 0.1% across the remaining windows.

#### Additional ECH analysis with masked archaic admixture

We observe an almost total depletion of archaic admixture in ECH haplotypes (see main text). However, archaic admixture will increase pairwise differences between the male haplotypes. If this reduces our ability to call ECHs overlapping admixed segments, it could explain why we only identify ECHs without admixture. To rule out this possibility, we repeated our inference procedure after masking all called admixture segments as missing data. The resulting calls of ECHs are thus unaffected by contributions to diversity by admixture. The result of this analysis is almost identical to the original analysis: Only an additional eighteen 100kb windows across chromosomes across samples are identified as ECHs. In comparison, the original analysis finds 13,808 such windows. ECHs are only called when five consecutive 100kb windows each contain more than 50,000 called bases. To allow calling of ECHs in the same set of windows after admixed segments are masked, we do not remove windows that fall below this cut-off after masking. However, in a small number of cases, admixture segments span a 100kb window, which means that a distance to other haplotypes cannot be computed after masking. 111 of these 100kb windows overlap haplotypes that are identified as ECH in other individuals. However, this only creates a very small uncertainty in the admixture proportion of ECH haplotypes: Even if we conservatively assume that all the 111 uncalled windows were, in fact, ECH haplotypes and only subsequently admixed, the admixture proportion of ECH haplotypes would only increase by a factor less than 1.01.

#### Haplotype structure plots

We follow (Crawford et al. 2017) to produce haplotype plots visualizing the relationship among haplotypes in the 90%-regions around each peak (given in Table S2). The left side of each figure is a UPGMA tree based on Jukes-Cantor corrected sequence distances between male haplotypes. The tree clusters the individual haplotypes, which are shown as horizontal lines on the right, color-coded according to geographical region. Vertical black bars on each haplotype represent non-reference SNPs. Plots for all regions are provided as supplementary data (Data S1).

#### Derived allele frequencies

We polarized all variants segregating in our sampled individuals using the chimpanzee (panTro3) variant provided in the Simons Genome Diversity Panel. Our analysis proceeds with X chromosomal SNPs in non-Africans, where we keep only SNPs called in at least 80% of non-Africans. This leaves 482,113 SNPs, each assigned to a 100kb window along the X chromosome. To characterize the relationship between non-African allele frequencies and ECH haplotypes, we compute across 100kb windows the proportion of derived variants with an allele frequency above 25%. Doing so separately for ECH and non-ECH windows shows the expected shift towards high-frequency alleles in ECH haplotypes (Figure 4).

#### ECHs shared with the ancient Ust’-Ishim male

The sequence of the Ust’-Ishim male is available as part of the Simons Genome Diversity data set. We computed pairwise sequence distances in 100kb windows between the Ust’-Ishim male and all present-day males. Windows, where the pairwise difference is based on fewer than 27,217 positions due to uncalled bases, are considered missing data. Ust’-Ishim has more uncalled bases than the contemporary samples, and the cutoff is adjusted from 50,000 to 27,217 to ensure the average number of 100kb pairwise comparisons to Ust’-Ishim matches the average number of 100kb pairwise comparisons between all contemporary samples. The distribution of these sequence distances to Ust’-Ishim shows the same characteristic enrichment of short pairwise distances as present-day non-Africans. To take the sampling time of the Ust'-Ishim into account when calling ECHs, we add a distance corresponding to 45,000 years when computing pairwise distances to present-day samples.

We repeated our clique-finding procedure to find ECHs, including the Ust'-Ishim male. The ECHs identified in the Ust'-Ishim male total 5.3 Mb. We find that the Ust'-Ishim shares the ECH with the contemporary non-Africans in six of the 14 most extreme regions (where the proportion of non-Africans sharing the ECH is at least 50%).

The very small number of sequence differences in ECHs implies that even a few base-calling errors will preclude inference of ECHs. For this reason, we performed a more robust and simpler inference of ECH sharing with Ust'-Ishim. We simply included the Ust'-Ishim in haplotype plots of each ECH. These plots are identical to those shown in Figure 5 but span only the of the 500kb centered at each peak in Figure 1A. This identifies additional three of the 14 most extreme regions where the Ust'-Ishim groups with the ECH haplotypes in contemporary non-Africans: 64,550,000-65,050,000; 76,800,000-77,300,000; 129,700,000-130,200,000. The ECH regions shared by the Ust'-Ishim are listed in Table S2.

### Estimation of the out-of-Africa bottleneck for SGDP samples

We used SMC++ on chromosome 7 to construct the autosomal population demography for the male samples analyzed. While the extensive population structure in the SGDP data set will inflate population sizes, this is not the case for the initial shared out-of-Africa bottlenecked population. The minimum Ne of this shared demography thus provides a good estimation of the bottleneck Ne relevant for the frequency simulations in Figure 7. In the SMC++ analysis, we use one male individual from each non-African region as the "dedicated individuals": S_Greek-2 (WestEurasia), S_Korean-1 (EastAsia), S_Irula-2 (SouthAsia), S_Yakut-2 (CentralAsiaSiberia), S_Papuan-9 (Oceania), S_Pima-1 (America). The remaining individuals contribute information though the SFS. We use 35 timepoints and estimate Ne for discrete epoques to better accommodate dramatic changes in population size. The demography estimated has a minimal bottleneck Ne of 4,126.

### Simulations

We use multinomial sampling to produce frequency trajectories of a population of haplotypes of length *L*, starting at frequency *1/N*. Each simulation is run for *G* generations. We generate n=100,000 such simulations. For the subset of simulations where any haplotype reaches the target frequency *f*, we identify the number of generations *g* until this happens. In each such case, we compute the probability that the haplotype did not recombine onto a different haplotype in the intervening *g* generations: $A_i = exp[-rLg] + (1 - exp[-rLg])/2$, where *r* is the recombination rate, and *g* is the generation in which frequency *f* is first reached. If *f* is not reached before generation *G*, then $A_i = 0$. The probability that a haplotype reaches frequency *f*, without recombining onto the background, is then estimated as the mean across sampled trajectories: $p = \frac{1}{n} \sum_i A_i$. The probability that at least one haplotype rises to frequency f along the entire chromosome is computed as: $1 - (1 - p)^{(S/W)}$, where *S* is the size of chromosome X (excluding the pseudoautosomal regions) and W is the window size of 500kb.

#### Analysis of CEU population in 1000 genomes data set

We use the phase3 version of the 1000 genome data set and extract the 49 males from the CEU population. We then repeat our ECH inference procedure using a clique size of 50% instead of 25%, which we used in the main analysis of SGDP. The larger clique size accommodates the additional recent ancestry among males from the homogeneous CEU population that does not occur between males sampled from the many separate populations in the SGDP. Using this clique size, the proportion of the X chromosome overlapping an ECH is 10%, similar to the 11% found in our SGDP analysis. Fifteen of the ECH peaks in the CEU analysis match exactly one of the nineteen peaks in the SGDP analysis (Figure S6). Four SGDP ECHs are not found in the CEU analysis, which in turn finds five ECHs not found in the SGDP analysis. The large overlap suggests that the majority of ECHs in the CEU sample arose in the early out-of-Africa population, shared by the SGDP populations.

#### Simulations of the CEU population from the 1000 genomes data set

Simulations represent the ancestral relationship of 49 male samples, mirroring the 49 male samples included in our analysis of ECHs in the CEU population. Among the two most detailed population demographic estimates of the CEU population previously published (Gravel et al. 2011; Terhorst et al. 2017), we chose the demography produced by (Gravel et al. 2011). Although the two are very similar in their representation, Gravel et al. report a marginally stronger bottleneck. We follow the approach by (Pool and Nielsen 2007) to scale the autosomal demography to reproduce the mean number of pairwise differences between CEU males on chromosome 7 (arbitrarily chosen for its intermediate size). We further apply two different scalings to represent the median X/autosome ratios of 0.51 (non-African samples) and 0.65 (African samples). For a realistic representation of recombination on the X chromosome, we use the sex-averaged deCODE recombination map derived from parent-offspring pairs (Kong et al. 2010). This map is not confounded by differences in effective population size or selection along the chromosome. We use SLiM3 (Haller and Messer 2019) to simulate fifteen consecutive 10Mb segments of the X chromosome from position 4,000,000 to 154,000,000, which excludes the pseudoautosomal regions. From each simulation, we extract 49 male X chromosomes (matching the number of CEU males) from the population and perform ECH inference as described above. On each simulated chromosome, we mask the same regions as missing data as we do in our analysis of the CEU population. We do this to ensure that the analyzed chromosomal regions from simulations correspond to the analyzed chromosomal regions in the CEU analysis.

#### Incomplete lineage sorting between human, chimpanzee, and gorilla

In a previous analysis (Munch et al. 2016), we estimated the proportion of incomplete lineage sorting (ILS) between humans, chimpanzees, and gorillas along the human genome. There, we analyzed autosomes and the X chromosome but published only estimates for autosomes. Extending this analysis, we computed the mean proportion of ILS in non-overlapping 1Mb windows along the X chromosome. Following (Munch et al. 2016), windows where the proportion of non-missing data falls below 30%, are not reported. The average proportion of ILS along the X chromosome is 13%. Following (Dutheil, Munch et al. 2015), mega-base windows are called as low-ILS windows if the proportion of ILS falls below 5%. These regions, totaling 12% of the chromosome, are shown as orange blocks in Figure 6. Jaccard tests are performed following (Favorov et al. 2012).

#### Gene ontology enrichment

To identify a list of candidate genes, we first identify the set of 100kb windows where the frequency of each ECH is maximal (Peak start and end in Table S2). We further include the immediately flanking 100kb windows to allow for the possibility that distant enhancers, rather than the transcribed region, is the target of selection on a gene. We tested the 65 protein-coding genes overlapping these regions for gene ontology enrichment using all protein-coding genes on the X chromosome as background. Performing this test against all gene ontology terms GOATOOLS (Klopfenstein et al. 2018) yielded no significant enrichments.

### Supplementary figures

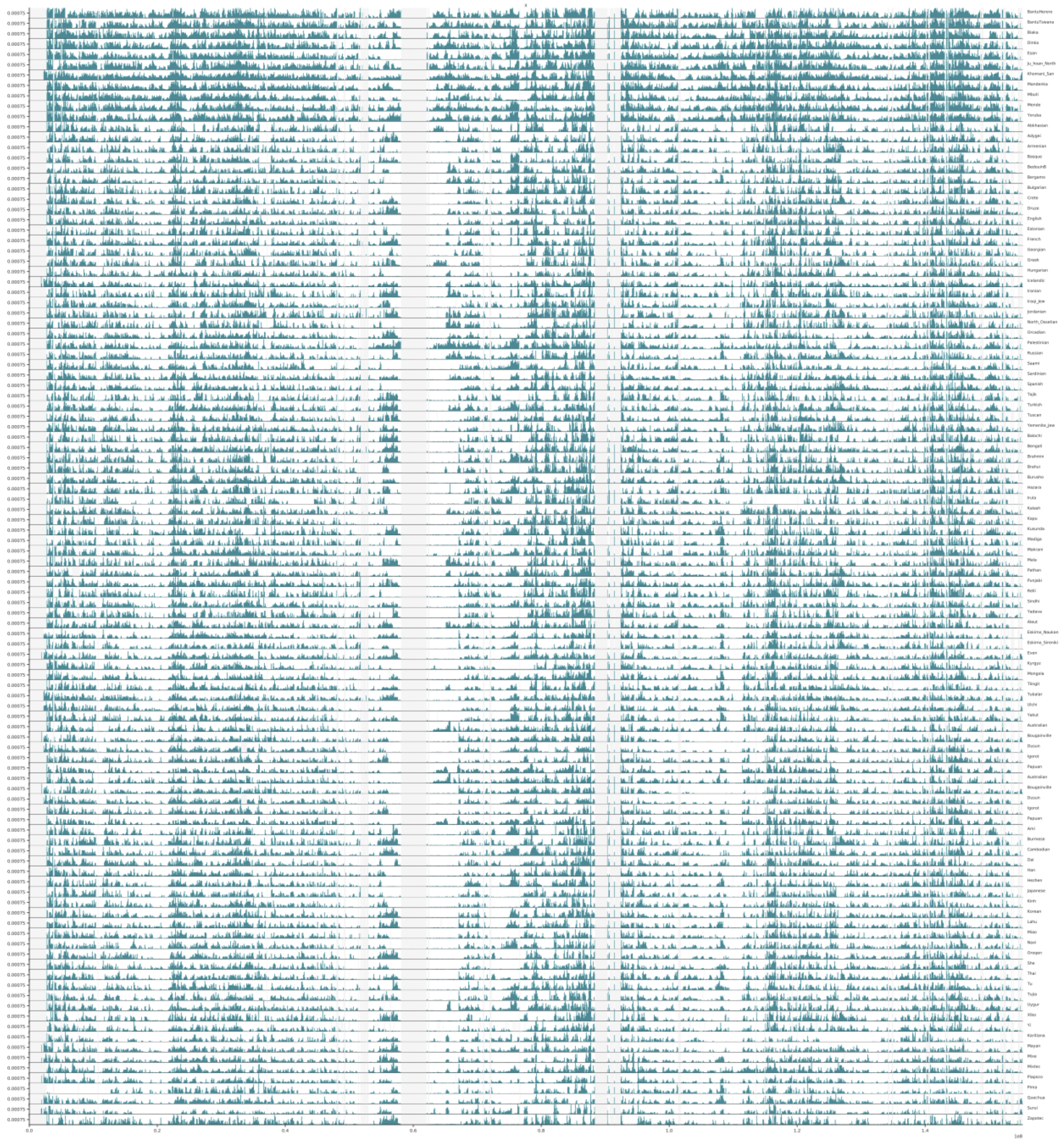

**Figure S1:** Mean pairwise differences in 100kb windows across chromosome X for each population. For a better visual comparison, the y-axis is truncated at 0.0015 to remove outliers. Light grey regions represent missing data.

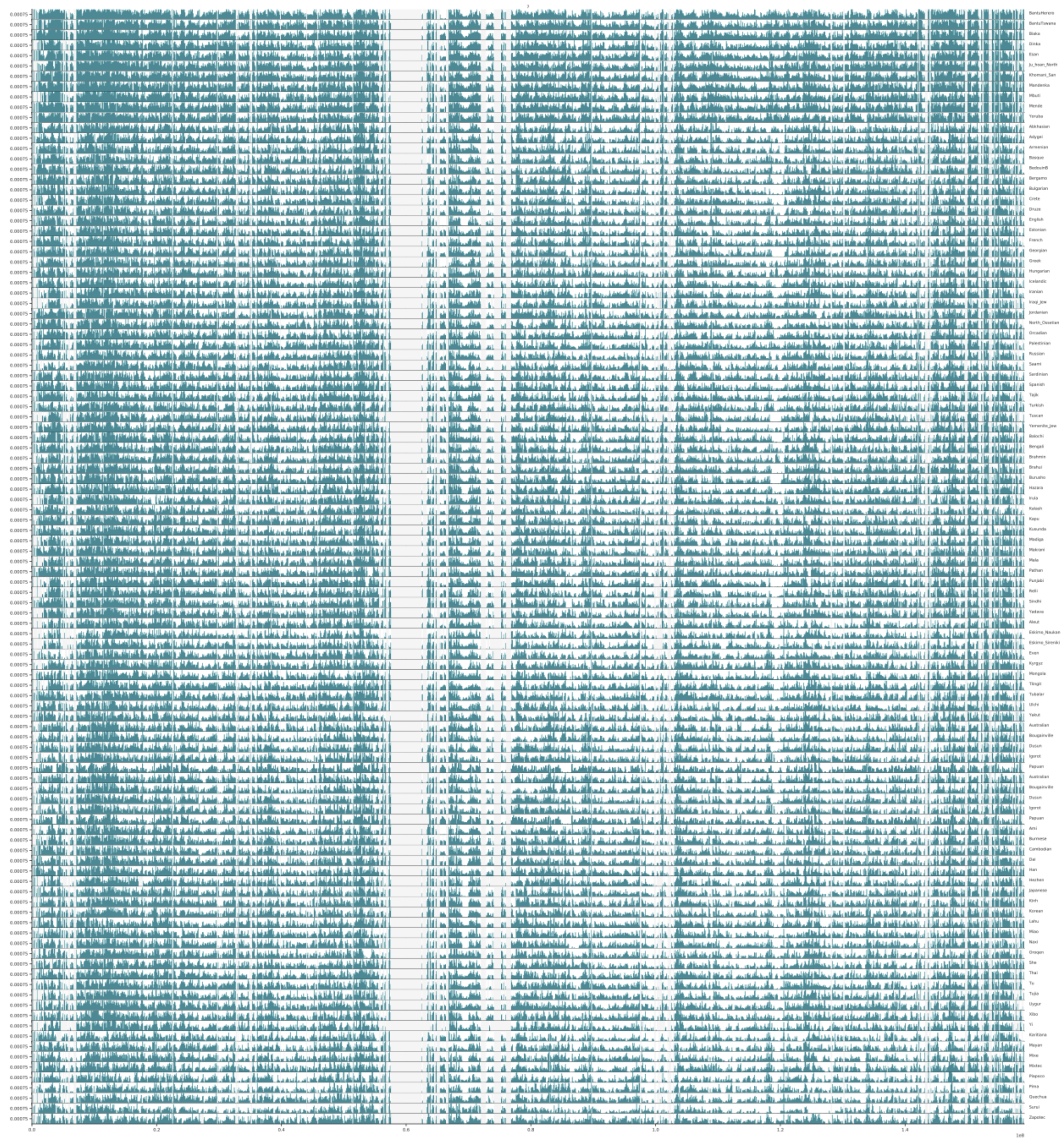

**Figure S2:** Mean pairwise differences in 100kb windows across chromosome 7 for each population. For a better visual comparison, the y-axis is truncated at 0.0015 to remove outliers. Grey regions represent missing data.

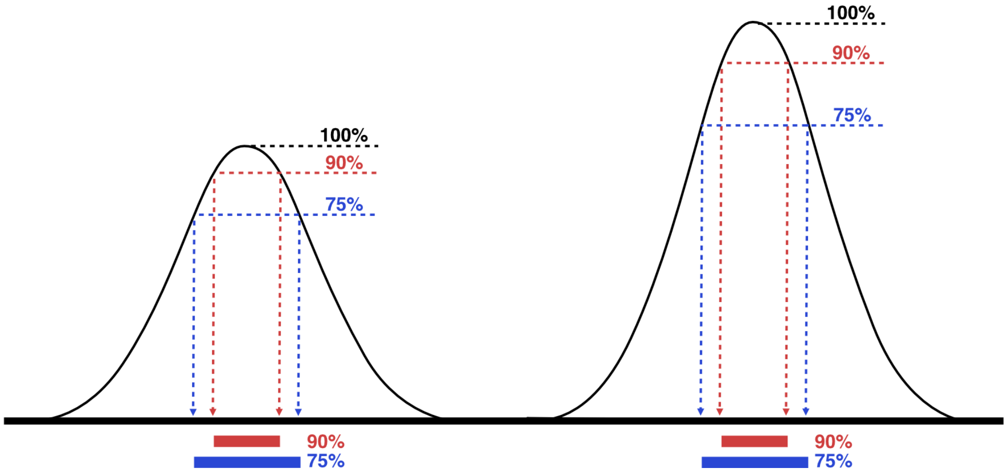


**Figure S3:** We operationally define a region around each peak where the proportion of individuals called as ECH is at least 90% of the peak value. Similarly, we define a wide region where the proportion of individuals called as ECH is above 75% of the peak value.

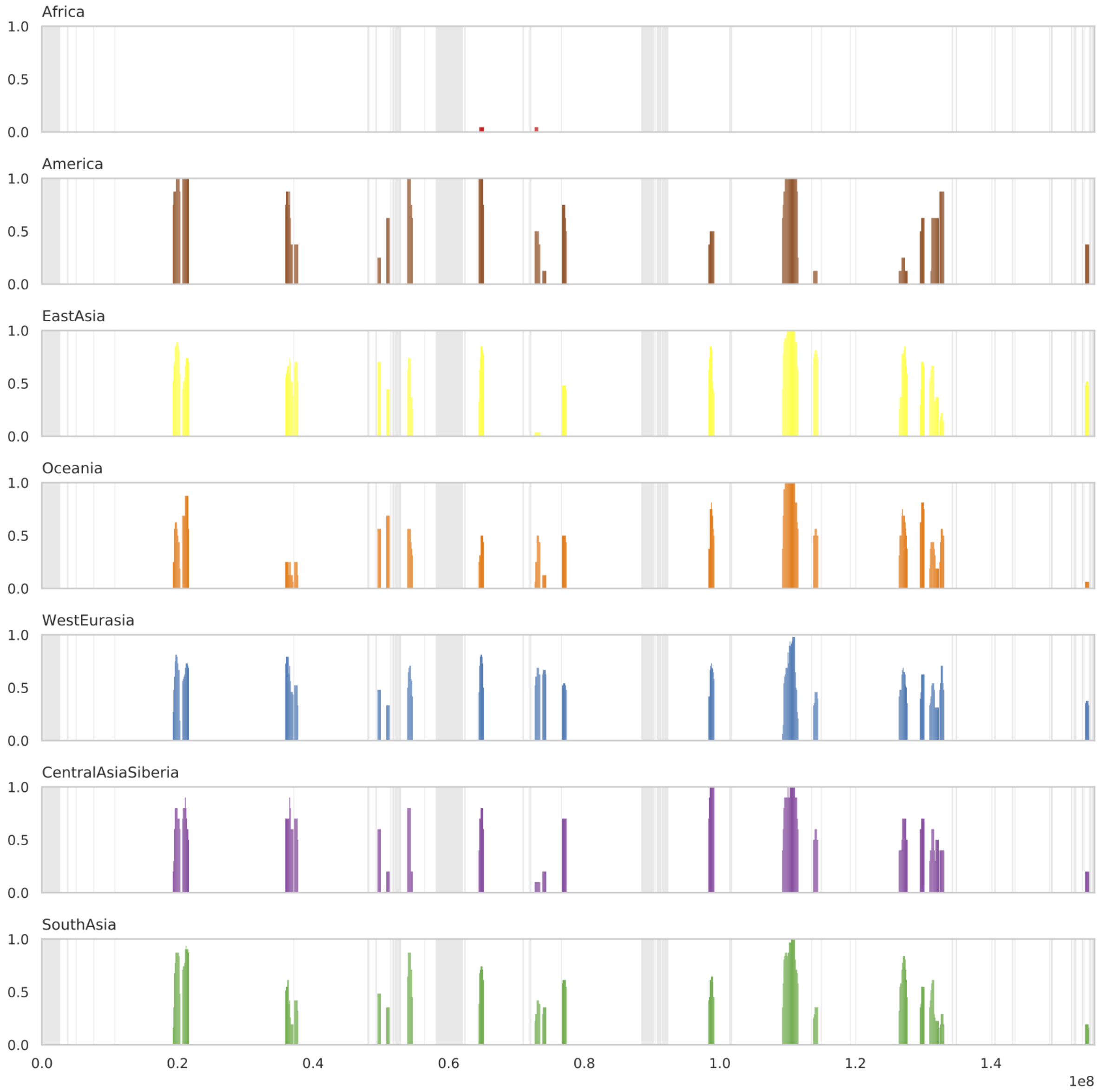


**Figure S4:** Proportion of individuals from each geographical region called as ECH in each 100kb window. Shaded regions represent regions with missing data.

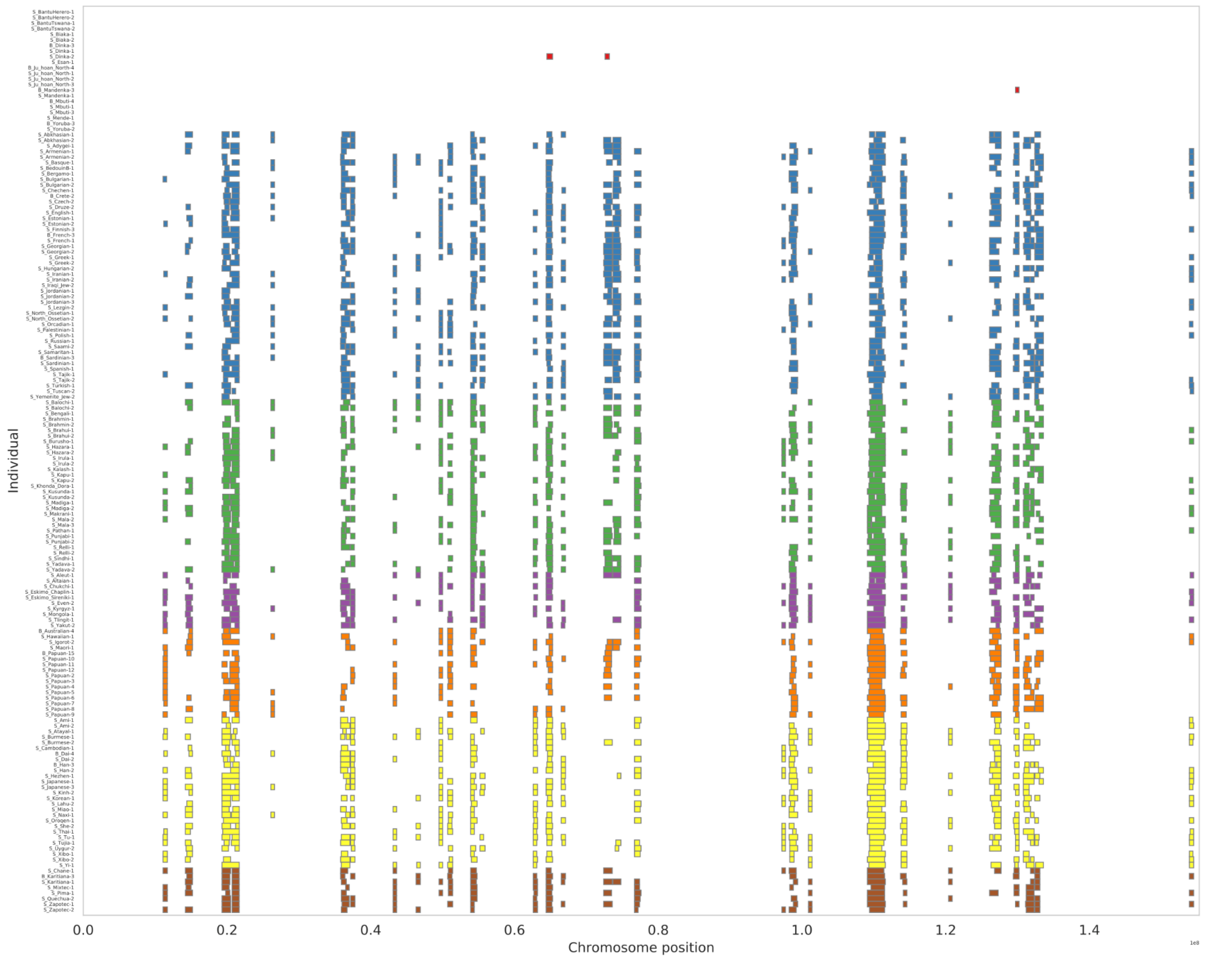


**Figure S5:** ECHs identified on each male X chromosome using 20% as the minimum number of individuals included in the haplotype clades defining an ECH.

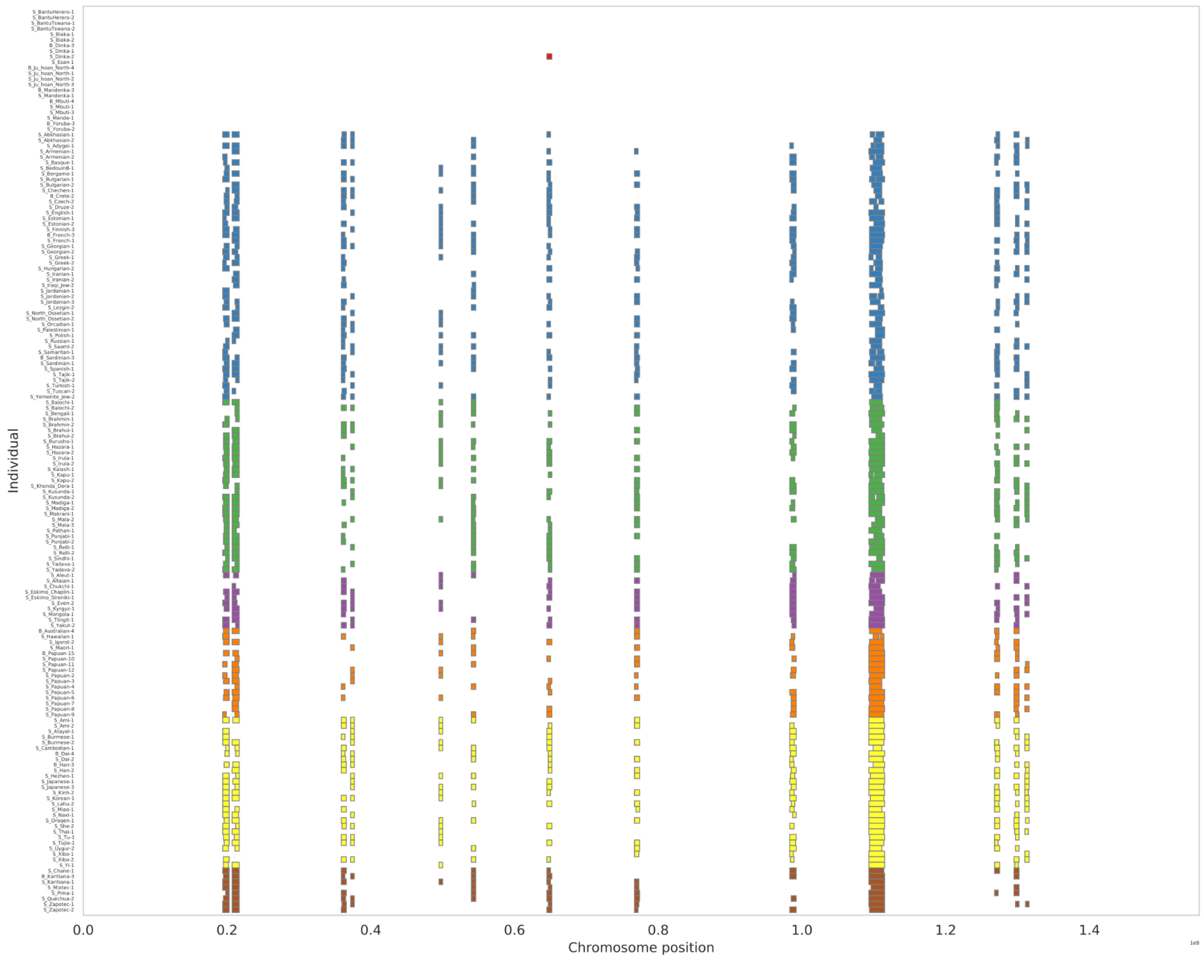


**Figure S6:** ECHs identified on each male X chromosome using 30% as the minimum number of individuals included in the haplotype clades defining an ECH.

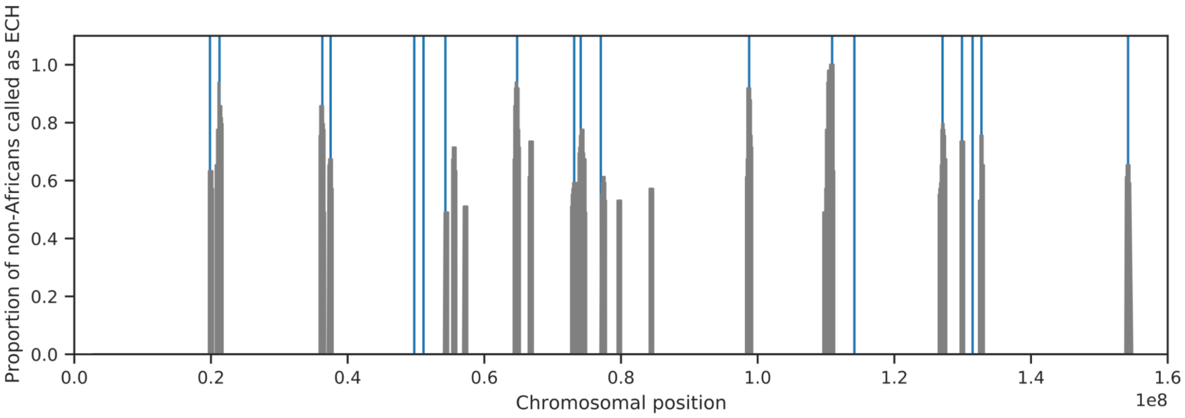


**Figure S7:** Frequency of ECHs along the X chromosome in the CEU population. Gray peaks show the proportion of 49 male haplotypes called as ECH in each 100kb window across the complete X chromosome. Vertical blue lines show the center of all 19 peaks identified in the SGDP analysis.

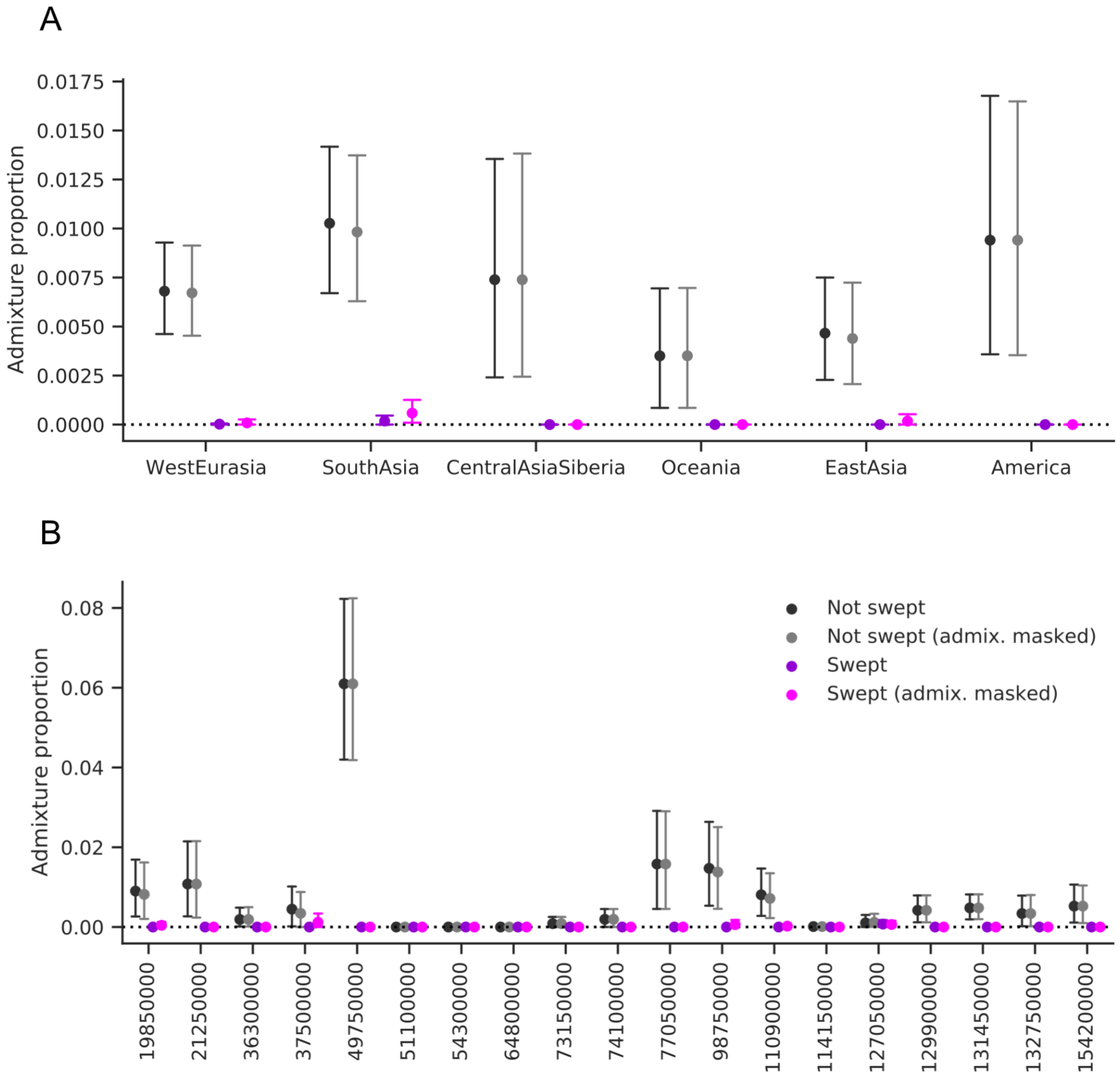


**Figure S8:** Admixture proportions on ECH and non-ECH haplotypes when masking inferred admixture before calling ECHs. Error bars with wide caps designate the standard 95% confidence intervals obtained from 10,000 bootstrapping iterations.

#### Supplementary Tables

**Table S1:** Male samples included in the analysis.

**Table S2:** Coordinates and statistics on each ECH.

**Table S3:** Admixture proportions when including only admixture segments sharing share derived variants with the high coverage archaic genomes (Denisova, Vindija, and Altai (DAV) segments).

#### Supplementary data

Supplementary data is available as “Data S1.zip”

#### References


Auton, A., Abecasis, G.R., Altshuler, D.M., Durbin, R.M., Abecasis, G.R., Bentley, D.R., Chakravarti, A., Clark, A.G., Donnelly, P., Eichler, E.E., et al. (2015). A global reference for human genetic variation. Nature *526*, 68–74. https://doi.org/10.1038/nature15393.

Bron, C., and Kerbosch, J. (1973). Algorithm 457: finding all cliques of an undirected graph. Commun Acm *16*, 575–577. https://doi.org/10.1145/362342.362367.

Crawford, N.G., Kelly, D.E., Hansen, M.E., Beltrame, M.H., Fan, S., Bowman, S.L., Jewett, E., Ranciaro, A., Thompson, S., Lo, Y., et al. (2017). Loci associated with skin pigmentation identified in African populations. Science *358*, eaan8433. https://doi.org/10.1126/science.aan8433.

Favorov, A., Mularoni, L., Cope, L.M., Medvedeva, Y., Mironov, A.A., Makeev, V.J., and Wheelan, S.J. (2012). Exploring Massive, Genome Scale Datasets with the GenometriCorr Package. PLoS Computational Biology https://doi.org/10.1371/journal.pcbi.1002529.

Klopfenstein, D.V., Zhang, L., Pedersen, B.S., Ramírez, F., Vesztrocy, A.W., Naldi, A., Mungall, C.J., Yunes, J.M., Botvinnik, O., Weigel, M., et al. (2018). GOATOOLS: A Python library for Gene Ontology analyses. Sci Rep-Uk 8, 10872. https://doi.org/10.1038/s41598-018-28948-z.

Lucotte, E.A., Skov, L., Jensen, J.M., Macià, M.C., Munch, K., and Schierup, M.H. (2018). Dynamic Copy Number Evolution of X- and Y-Linked Ampliconic Genes in Human Populations. Genetics *209*, 907–920. https://doi.org/10.1534/genetics.118.300826.

Mallick, S., Li, H., Lipson, M., Mathieson, I., Gymrek, M., Racimo, F., Zhao, M., Chennagiri, N., Nordenfelt, S., Tandon, A., et al. (2016). The Simons Genome Diversity Project: 300 genomes from 142 diverse populations. Nature *538*. https://doi.org/10.1038/nature18964.

Munch, K., Nam, K., Schierup, M., and Mailund, T. (2016). Selective Sweeps across Twenty Millions Years of Primate Evolution. Molecular Biology and Evolution 33, msw199. https://doi.org/10.1093/molbev/msw199.

Sankararaman, S., Mallick, S., Patterson, N., and Reich, D. (2016). The Combined Landscape of Denisovan and Neanderthal Ancestry in Present-Day Humans. Current Biology 26, 1241–1247. https://doi.org/10.1016/j.cub.2016.03.037.

Skov, L., Hui, R., Shchur, V., Hobolth, A., Scally, A., Schierup, M., and Durbin, R. (2018). Detecting archaic introgression using an unadmixed outgroup. Plos Genet 14, e1007641. https://doi.org/10.1371/journal.pgen.1007641.

Auton A., G. R. Abecasis, D. M. Altshuler, R. M. Durbin, G. R. Abecasis, *et al.*, 2015 A global reference for human genetic variation. Nature 526: 68–74. https://doi.org/10.1038/nature15393

Bron C., and J. Kerbosch, 1973 Algorithm 457: finding all cliques of an undirected graph. Commun Acm 16: 575–577. https://doi.org/10.1145/362342.362367

Crawford N. G., D. E. Kelly, M. E. Hansen, M. H. Beltrame, S. Fan, et al., 2017 Loci associated with skin pigmentation identified in African populations. Science 358: eaan8433. https://doi.org/10.1126/science.aan8433

Favorov A., L. Mularoni, L. M. Cope, Y. Medvedeva, A. A. Mironov, et al., 2012 Exploring Massive, Genome Scale Datasets with the GenometriCorr Package. PLoS Computational Biology. https://doi.org/10.1371/journal.pcbi.1002529
Fu Q., H. Li, P. Moorjani, F. Jay, S. M. Slepchenko, et al., 2014 Genome sequence of a 45,000-year-old modern human from western Siberia. Nature 514: 445–449. https://doi.org/10.1038/nature13810
Gravel S., B. M. Henn, R. N. Gutenkunst, A. R. Indap, G. T. Marth, et al., 2011 Demographic history and rare allele sharing among human populations. Proc National Acad Sci 108: 11983–11988. https://doi.org/10.1073/pnas.1019276108
Haller B. C., and P. W. Messer, 2019 SLiM 3: Forward Genetic Simulations Beyond the Wright–Fisher Model. Mol Biol Evol 36: 632–637. https://doi.org/10.1093/molbev/msy228
Klopfenstein D. V., L. Zhang, B. S. Pedersen, F. Ramírez, A. W. Vesztrocy, et al., 2018 GOATOOLS: A Python library for Gene Ontology analyses. Sci Rep-uk 8: 10872. https://doi.org/10.1038/s41598-018-28948-z
Kong A., G. Thorleifsson, D. F. Gudbjartsson, G. Masson, A. Sigurdsson, et al., 2010 Fine-scale recombination rate differences between sexes, populations and individuals. Nature 467. https://doi.org/10.1038/nature09525
Lucotte E. A., L. Skov, J. M. Jensen, M. C. Macià, K. Munch, et al., 2018 Dynamic Copy Number Evolution of X- and Y-Linked Ampliconic Genes in Human Populations. Genetics 209: 907–920. https://doi.org/10.1534/genetics.118.300826
Mallick S., H. Li, M. Lipson, I. Mathieson, M. Gymrek, et al., 2016 The Simons Genome Diversity Project: 300 genomes from 142 diverse populations. Nature 538. https://doi.org/10.1038/nature18964
Munch K., K. Nam, M. Schierup, and T. Mailund, 2016 Selective Sweeps across Twenty Millions Years of Primate Evolution. Molecular biology and evolution 33: msw199. https://doi.org/10.1093/molbev/msw199
Pool J. E., and R. Nielsen, 2007 POPULATION SIZE CHANGES RESHAPE GENOMIC PATTERNS OF DIVERSITY. Evolution 61: 3001–3006. https://doi.org/10.1111/j.1558-5646.2007.00238.x
Sankararaman S., S. Mallick, N. Patterson, and D. Reich, 2016 The Combined Landscape of Denisovan and Neanderthal Ancestry in Present-Day Humans. Current Biology 26: 1241–1247. https://doi.org/10.1016/j.cub.2016.03.037
Skov L., R. Hui, V. Shchur, A. Hobolth, A. Scally, et al., 2018 Detecting archaic introgression using an unadmixed outgroup. Plos Genet 14: e1007641. https://doi.org/10.1371/journal.pgen.1007641
Terhorst J., J. A. Kamm, and Y. S. Song, 2017 Robust and scalable inference of population history from hundreds of unphased whole genomes. Nature Genetics 49: 303–309. https://doi.org/10.1038/ng.3748
